# A nonlinear time-domain finite-element model of the human middle ear subjected to intense sound waves

**DOI:** 10.64898/2026.09.23.753568

**Authors:** Hossein Mohammadi, Andrew Tubelli, Sunil Puria, Jeffrey Tao Cheng

## Abstract

Computational models of the middle ear are typically developed within the linear range of the middle ear’s response, which corresponds to typical sounds in daily life. Understanding middle-ear biomechanics under high-intensity sounds is also important for assessing potential auditory damage under adverse conditions. In this study, a finite-element (FE) model of the human middle ear plus the external ear canal is presented, employing a first-order Ogden hyperelastic material formulation and time-domain nonlinear analysis to capture nonlinear deformation of soft tissues and the resulting middle-ear response. The model was analyzed using pure-tone pressure excitations applied at the ear canal entrance over the 80 Hz to 8 kHz frequency range and 110 to 180 dB sound pressure levels (SPL). The ear canal pressure near the tympanic membrane (TM) and full-field three-dimensional response of the middle ear structures were examined. Across the investigated frequency range, the simulated middle-ear motion exhibited a transition from predominantly piston-like behavior at low frequencies, up to 1 kHz, to increasingly intricate multi-modal and multi-directional vibration patterns at higher frequencies. Nonlinear behaviors became increasingly significant at high frequencies and high stimulus levels, including compressive/expansive pressure-displacement relationships, waveform distortions, and the emergence of non-harmonic spectral components. The model developed in this study provides a computational framework for studying nonlinear sound transmission through the middle ear and its implications for high-intensity acoustic exposure. The findings contribute to a better understanding of middle-ear mechanics under adverse conditions and support future investigations of acoustic trauma and hearing protection strategies.

## 1. Introduction

The human middle ear is generally considered to behave as a linear system, as it conducts low- to moderate-level sounds into mechanical vibrations transmitted to the inner ear. However, under high-intensity sounds, usually above 130 dB sound pressure level (SPL)^1^, nonlinear responses of the middle ear have been observed in anesthetized animals or cadaveric temporal bones without middle ear muscle reflex, shown as either compressive or expansive growth of middle-ear displacement with respect to sound pressure excitation (Guinan and Peake 1967; Rosowski et al. 2019; Cheng et al. 2021). Despite the existence of such “passive” nonlinear behaviors, the underlying mechanical mechanisms responsible for them remain poorly understood. Computational approaches provide a way to analyze the mechanical behavior of middle-ear structures under varying acoustic loads and material settings to better understand these nonlinear responses.

The finite-element (FE) method, a powerful computational tool, has been extensively used for *in silico* modelling of middle-ear mechanics, starting from the pioneering work of (Funnell and Laszlo 1978) to more recent and advanced models reported in (Thompson et al. 2023; Ebrahimian et al. 2024a, 2025; Nørgaard et al. 2024; Muyshondt et al. 2026). With the development of computational resources over the past decade, more complicated geometries, boundary conditions, and material properties are incorporated with the FE approach, sometimes combined with machine learning and surrogate modelling techniques for faster response approximation (Ebrahimian et al. 2023a, 2024b; Motallebzadeh et al. 2024, 2025; Asakura et al. 2026). However, previous FE middle-ear models have mainly focused on the linear regime without introducing nonlinear effects in geometry and mechanical properties.

The ossicles, as the bony structures of the middle ear, can be reliably defined using linear elastic material models, even under high-intensity sounds, as they do not undergo large strain-induced deformation.

However, soft tissues of the middle ear, including the tympanic membrane (TM), joints, ligaments, and tendons, are usually susceptible to large-strain deformation, and linear material models are not sufficiently accurate at high excitation levels. Researchers have conducted a limited number of studies on the nonlinear properties of middle-ear soft tissues. Ladak et al. (2006) developed a model of the cat TM that incorporated geometrical nonlinearity while assuming a linear constitutive law for the material model. Others have used nonlinear models for isolated tissue corresponding to individual structures of the human middle ear. These include the stapedial tendon (ST) (Cheng and Gan 2007), tensor tympani tendon (TTT) (Cheng and Gan 2008a), anterior malleal ligament (AML) (Cheng and Gan 2008b), stapedial annular ligament (SAL) (Gan et al. 2011), incudostapedial joint (ISJ) (Zhang and Gan 2011), and TM (Cheng et al. 2007; Motallebzadeh et al. 2013), by implementing the Ogden hyperelastic model into FE simulations. Liang et al. (2016) also performed an inverse FE analysis on chinchilla TM specimens and obtained second-order Ogden models under static loading conditions.

However, developing a complete nonlinear model of the middle ear is more challenging, as it is computationally expensive and prone to numerical convergence issues. It is also difficult to optimize such models because of the larger number of parameters involved and longer simulation times required to fit experimental data. Wang et al. (2007) developed a nonlinear FE model to analyze the static response of the middle ear by introducing both geometric and material nonlinearity into the TM, ligaments, and muscle tendons using a five-parameter Mooney-Rivlin hyperelastic model. They also analyzed the dynamic behavior of the middle ear; however, linear elastic properties based on a stress-dependent modulus were used in those simulations. Additionally, Qi et al. (2008) implemented the Mooney-Rivlin model into the TM, AML, and posterior incudal ligament (PIL) in a static nonlinear model of the middle ear. More recently, Mohammadi et al. (2024a, b; 2026) incorporated nonlinear properties into the TM using an Ogden hyperelastic material model while modelling the deformation of the entire middle ear under quasi-static needle insertion, which imposed large-strain deformation on the TM. They showed that a first-order Ogden hyperelastic model can accurately capture the mechanical nonlinearity of the TM under large-strain quasi-static puncture loading.

Muyshondt and Dirckx (2021) used the Veronda–Westmann hyperelastic model to account for the nonlinear elastic deformation of soft tissues in the human middle ear under large static pressure (up to 4 kPa). Moreover, they performed frequency-domain dynamic analysis using a linearized hyperelastic model around a preloaded state (i.e., external static pressure) to solve harmonic vibrations under 1 Pa dynamic excitation. Consequently, they developed new models of the rabbit middle ear to analyze the effects of prestrain in the TM on hearing response using the same modelling framework, i.e., hyperelastic static analysis and linearized harmonic response around a prestressed state (Muyshondt and Dirckx 2022). However, they used the Veronda–Westmann model only for the *pars tensa*, SAL, and joints, while a simpler Neo-Hookean model was used for the PIL and *pars flaccida*. They demonstrated that changes in geometry, prestress or prestrain, and material nonlinearity all contribute to pressure-induced stiffening and alterations in the middle ear’s response.

In general, the Ogden hyperelastic material model has been widely used as a constitutive model for analyzing soft tissue biomechanics, including the middle ear (Motallebzadeh et al. 2013; Liang et al. 2016; Lohr et al. 2022; Mohammadi et al. 2024b). Additionally, experimentally fitted ranges of mechanical properties for the Ogden model exist for soft tissues extracted from the middle ear, enabling the development of a complete nonlinear model of the middle ear. In this study, we present a dynamic nonlinear model of the middle ear by introducing nonlinearity into all middle-ear soft tissues using the Ogden hyperelastic material formulation while modelling its response to high-intensity pure-tone sinewave excitation in the time domain. This approach allowed us to analyze both the transient and steady-state responses of the middle ear at different excitation levels and frequencies.

## 2. Materials and Methods

The methodology consisted of conducting FE modelling simulations with data analysis as follows.

### 2.1. Finite-Element Model

This section describes the FE model, including the model configuration, governing equations and boundary conditions, material properties, and numerical implementation.

#### 2.1.1. Model Configuration

We used a previously developed FE model of the middle ear and ear canal reconstructed from high-resolution μCT images of a human temporal bone (Motallebzadeh et al. 2025). Figure 1 illustrates the FE model from the posterior (Fig. 1a) and medial (Fig. 1b) views. The model included a short section of the ear canal (approximately 6.6 mm from the umbo to the center of the input plane), modelled as an acoustic domain, and middle-ear structures modelled as solid domains. A common anatomical coordinate system was used in the model. A lateral-medial (LM) direction was defined to be approximately normal to the stapes footplate (SFP), such that the LM component closely represents the piston-like motion of the stapes. The two other components represent approximate anterior-posterior (AP) and superior-inferior (SI) directions. Since the SFP is not perfectly planar and the TM is oriented differently from the SFP, the LM direction does not exactly coincide with the local surface normal at every location. Therefore, the AP, SI, and LM components should be interpreted as global anatomical directions rather than local surface-based directions. This common coordinate system facilitates direct comparison of kinematics between the TM and SFP.

**Fig. 1.**
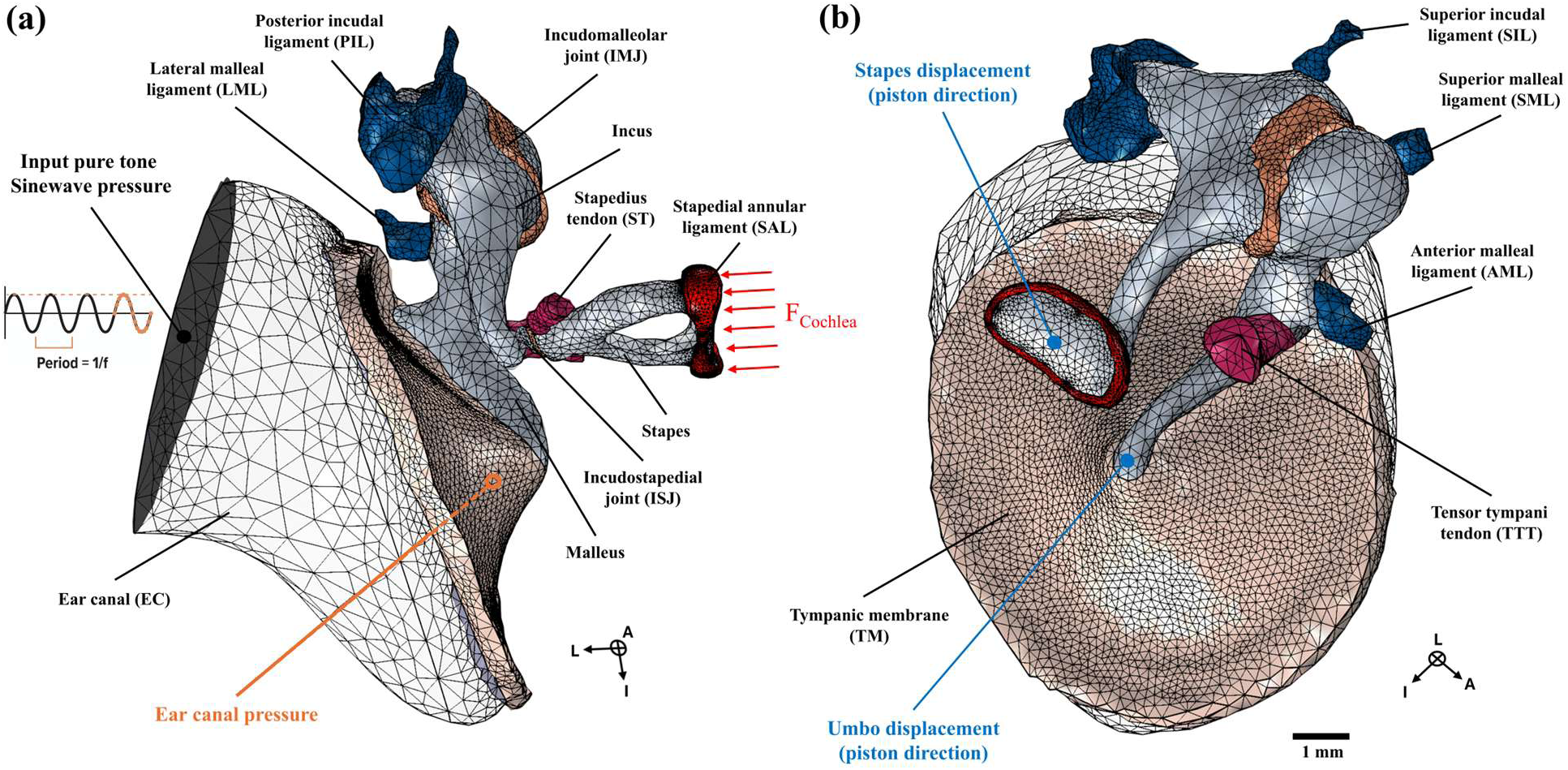
Meshed FE model of the ear canal and middle-ear structures, including the ossicles and soft tissues (i.e., TM, ligaments, joints, and tendons), visualized from the posterior side (a) and medial side (b). The Cartesian coordinate system approximately indicates the lateral (L), anterior (A), and inferior (I) directions. Pure-tone sinusoidal pressure was applied to the entire surface of the entrance of the shortened ear canal as the model input. The cochlear load (F_Cochlea_) was represented by a viscous impedance, shown as a distributed load applied to the SFP. Ear canal pressure (1 mm away from the umbo), 3D umbo displacement, and 3D stapes displacement (at the center of the footplate) were measured as model outputs. A right-handed coordinate system (I-L-A) is shown in both panels

The middle ear consisted of the malleus, incus, stapes, TM, incudomalleolar joint (IMJ), ISJ, and multiple suspensory ligaments and tendons, including the superior malleal ligament (SML), superior incudal ligament (SIL), lateral malleal ligament (LML), AML, PIL, SAL, TTT, and ST. The FE model consisted of 74,041 tetrahedral elements with a total mesh volume of 303.7 mm^3^. Quadratic elements were used in both the acoustic and solid domains to improve the accuracy of model predictions, considering the geometrical complexities and large-strain deformations under high-intensity sounds. The solid domains were discretized using quadratic serendipity elements to improve computational efficiency, reduce memory usage, and reduce solution times. The middle-ear cavity was excluded due to its expected linear nature and its impact on computational efficiency.

#### 2.1.2. Governing Equations and Boundary Conditions

The ear canal acoustic domain was modelled using the lossless wave equation as follows:

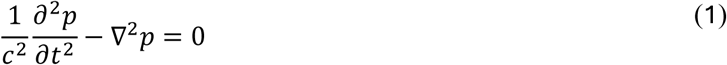

where *t* represents time, *p* is the acoustic pressure, *c* is the speed of sound, 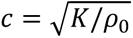, *K* is the bulk modulus, and *ρ*_0_ is the density of the air. The air was modelled at a default temperature of 20 ^◦^C and an absolute pressure of 1 atm. The transient pressure of the acoustic model was defined using a linear elastic fluid model, with a density, *ρ_0_*, of 1.2 kg/m^3^ and speed of sound, *c*, equal to 343.2 m/s.

The rest of the middle ear, comprising the structural domain, was governed by the momentum conservation equation defined by:

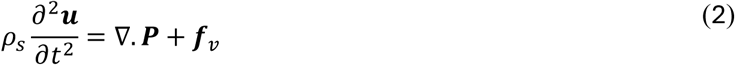

where *ρ_s_* represents solid density, ***u*** is the displacement vector, ***P*** is the first Piola-Kirchhoff stress tensor in the reference configuration, and ***f_v_*** is the body force.

The acoustic and structural domains were coupled at the TM interface through continuity of normal velocity and force equilibrium. The normal acceleration of the fluid was enforced to match the structural acceleration at the interface, ensuring kinematic compatibility. In addition, the acoustic pressure acted as a normal traction on the structure, such that:

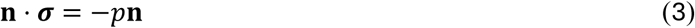

where **n** is the unit normal vector and ***σ*** is the Cauchy stress tensor. These conditions establish a two-way coupling between the acoustic pressure field and structural vibrations.

On the acoustic part of the model, a rigid-wall boundary condition with zero normal particle velocity was used on the ear canal walls. Loading was defined using pure-tone sinewave excitation at the entire entrance surface of the ear canal in the model. On the solid part of the model, the outer surfaces of the TM annulus and the SAL, as well as the outer ends of the ligaments (i.e., SIL, PIL, AML, SML, and LML) and tendons (i.e., TTT and ST), were fixed using zero-displacement boundary conditions based on literature (Maftoon et al. 2015; O’Connor et al. 2017; Ebrahimian et al. 2023b).

The cochlear fluid load was modelled using an acoustic input impedance, *Z*_c_, of 20 GΩ (Gan et al. 2004a), which lies within the reported range for human cochlear impedance in the low- to mid-frequency region (Merchant et al. 1996; Aibara et al. 2001). The impedance was defined as the pressure-to-volume velocity ratio at the SFP and implemented as an equivalent surface damping coefficient, *C_vd_* = *Z_c_A_fp_* = 5.99 × 10^4^ Pa ⋅ s/m, using a footplate area of *A_fp_* = 2.99 mm^2^. The mass and stiffness contributions of the cochlear load were neglected for simplicity (Puria and Allen 1991; Shera and Zweig 1991).

To account for geometrical nonlinearity in the model, the Green-Lagrange strain tensor was used:

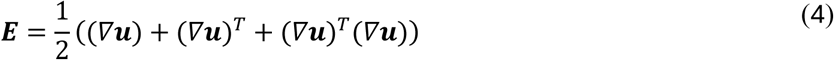

where ***E*** denotes the Green–Lagrange strain tensor, ***u*** is the displacement vector field, and *∇**u*** is the displacement gradient tensor with respect to the reference configuration. Superscript *T* denotes the transpose operator.

The simulations were conducted in the time domain with tone inputs at discrete frequencies, including 80, 250, 500, 1000, 2000, 4000, and 8000 Hz, and levels of 110, 120, 130, 140, 150, 160, 170, and 180 dB SPL (re 20 µPa). Excitation durations were 16 ms consistently across all investigated cases except for the lowest frequency of 80 Hz, where a duration of 50 ms was used. This resulted in 4 cycles at 80 and 250 Hz, and 8, 16, 32, 64, and 128 cycles at 500, 1000, 2000, 4000, and 8000 Hz, respectively. Fewer cycles were included at lower frequencies where more steady-state behavior was expected, whereas more cycles were included at higher frequencies where more transient behavior was expected. At high frequencies, fewer pressure levels were included due to numerical convergence issues in high-level, high-frequency cases. For instance, only levels up to 150 dB SPL successfully converged at 4 kHz and 8 kHz.

#### 2.1.3. Material Properties

The densities of malleus, incus, and stapes were 2390 kg/m^3^, 2150 kg/m^3^, and 2200 kg/m^3^, respectively, while a density of 1100 kg/m^3^ was used for all soft tissues (Sim and Puria 2008; Maftoon et al. 2015; O’Connor et al. 2017). Since the ossicles are relatively stiff and do not undergo large-strain deformation, exhibiting maximum first principal strains below 0.6% even under the highest sound levels considered in this study, linear elastic material models can be reliably used to represent their mechanical behavior (Mohammadi et al. 2026). Therefore, the ossicles were modeled as linear elastic materials with a Young’s modulus of 14 GPa and a Poisson’s ratio of 0.3 (O’Connor et al. 2017; Ebrahimian et al. 2023b). In contrast, a first-order Ogden hyperelastic material model (Charlebois et al. 2013; Motallebzadeh et al. 2013) was used for the soft tissues to account for their nonlinear behavior under high-intensity sounds (Ogden 1972):

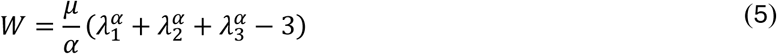

where *λ_i_* (*i* = 1, 2, 3) are the principal stretches, and *μ* and *α* are the Ogden constants. The parameter *μ* primarily affects the overall stiffness of the tissue, whereas *α* controls the curvature of the stress–strain response and therefore the extent of nonlinear behavior at large deformations. Table 1 provides the Ogden constants that we used for all included middle-ear soft tissues in our FE simulations. These values represent mean experimental values reported in the literature. It is worth noting that the experimental constants are based on a modified form of the Ogden equation, where *μ* represents the shear modulus of each soft tissue.^2^

**Table 1:**
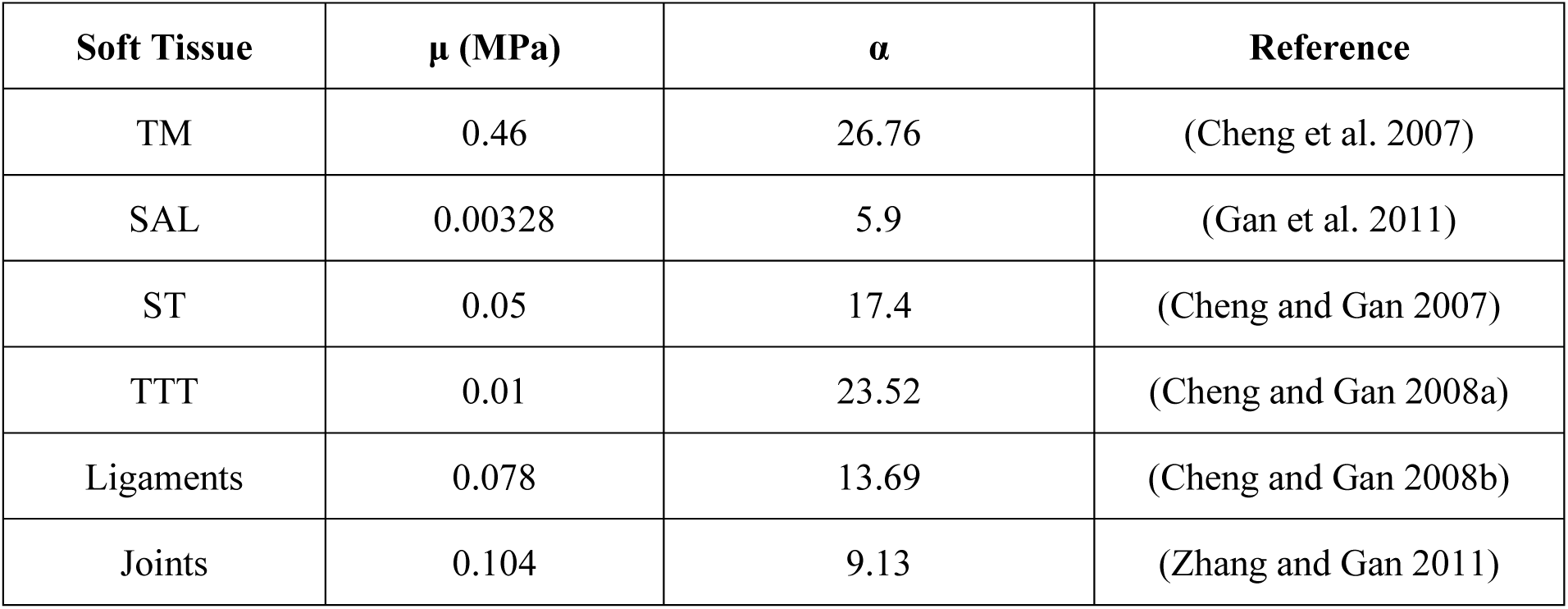
First-order Ogden model parameters for soft tissues included in the FE model of the middle ear.

| Soft Tissue | $\mu$ (MPa) | $\alpha$ | Reference |
| --- | --- | --- | --- |
| TM | 0.46 | 26.76 | (Cheng et al. 2007) |
| SAL | 0.00328 | 5.9 | (Gan et al. 2011) |
| ST | 0.05 | 17.4 | (Cheng and Gan 2007) |
| TTT | 0.01 | 23.52 | (Cheng and Gan 2008a) |
| Ligaments | 0.078 | 13.69 | (Cheng and Gan 2008b) |
| Joints | 0.104 | 9.13 | (Zhang and Gan 2011) |

#### 2.1.4. Numerical Implementation

All models were developed and simulated in COMSOL Multiphysics®, version 6.4 (COMSOL Multiphysics, Burlington, MA, USA). Simulations were performed in the time domain using an implicit solver with the generalized-α method. A convergence study on the generalized-α method showed that a time step resolution of 60 per cycle [1/(60*f_excitation_*)] achieved almost perfect convergence at low to mid frequencies (i.e., up to 2 kHz) and reliable convergence at higher frequencies (i.e., 4 kHz and 8 kHz). All simulations up to 1 kHz were conducted at a higher time stepping resolution of 1/(60,000), as they were relatively fast, while a time stepping of 1/(60*f_excitation_*) was used for 2, 4, and 8 kHz, for computational efficiency reasons. A constant Newton approach was used for the nonlinear solver configuration. Simulations were performed on a 40-core workstation (Intel Xeon E5-2698 v4, 512 GB RAM) using the PARDISO solver, selected for its superior shared-memory speedup and efficiency on high-core-count systems (Schenk and Gärtner 2002). Additionally, the transient response was sampled using a fixed number of output points corresponding to 16 samples per excitation cycle for each frequency, which is well above the Nyquist sampling requirement (*f_s_* ≥ 2*f_max_*) (Jerri 1977). This ensured adequate temporal resolution for accurately capturing the waveform and steady-state behavior. We performed 46 separate simulations covering all the investigated frequencies and SPLs, with a total computational time of approximately 358 hours. Computational time generally increased with both level and frequency. The lowest computational times were observed for frequencies below 1 kHz, averaging approximately 3–4 hours per simulation for a 16 ms time-domain analysis, whereas the highest computational cost occurred at 8 kHz, where each simulation required approximately 22 hours for the same 16 ms analysis.

### 2.2. Data Analysis

This section describes the data analysis procedures applied to the FE time-domain results, including full-field and transient analyses, steady-state determination, spectral analysis, and incremental nonlinearity index analysis.

#### 2.2.1. Full-field and Transient Analyses

FE simulations provided time-domain three-dimensional displacement of the entire middle ear and ear canal pressure over time. We analyzed the full-field displacement of the middle-ear structures at a peak instance in the last cycle of the simulation based on the input pressure signal at multiple representative frequencies, including 0.5, 4, and 8 kHz, each at a lower and a higher excitation level. For this analysis, we used the magnitude of displacements. Additionally, we selectively analyzed the ear canal pressure at 1 mm from the umbo, and the displacements at the umbo and center of the SFP in the time domain for the same representative frequencies and levels. For the transient analysis, all three orthogonal motion components, including AP, SI, and LM directions, were taken into consideration.

#### 2.2.2. Steady-State Determination

The simulated responses exhibited an initial transient phase before reaching steady-state behavior (see time-domain results in Section 3.3). Therefore, an objective procedure was used to identify the onset of steady state so that subsequent analyses were performed only on the steady-state portion of each signal. A delay-coordinate embedding approach was used to construct Poincaré representations (Shahhosseini et al. 2023) of the displacement signals to assess the attainment of steady-state behavior. For a given time-domain signal *u*(*t*), the embedded coordinates were defined as:

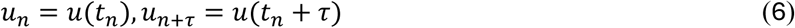

where *τ* is a fixed time delay, which was selected as a fraction of the excitation period, specifically *τ = T/8*, where *T = 1/f_exc_* is the excitation period. This choice ensured that the embedded coordinates captured the temporal evolution of the oscillatory response while maintaining sufficient separation between correlated samples. The resulting trajectory in the (*u*(*t*), *u*(*t* + *τ*)) plane provided a reduced phase-space representation of the system dynamics and enabled visualization of periodicity and deviations from a pure-tone waveform.

To quantitatively determine the duration required to reach a steady-state response, the time-domain signals for the umbo and SFP displacements were divided into equal-length temporal segments. The normalized difference between consecutive segments was defined using the Euclidean norm:

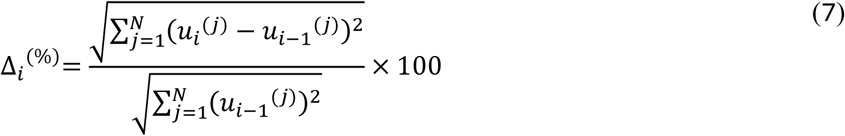

where 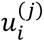 denotes the *j*-th sample of the response in segment *i*, and *N* is the number of samples in each segment. The quantity 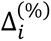 corresponds to the normalized percentage change between two consecutive segments. For all frequencies, the time-domain signals were divided into eight segments, except for 80 Hz and 250 Hz, where the simulated signal contained only four cycles and the response was therefore divided into four segments. In all cases, the signals were sampled at 16 points per cycle, resulting in fewer samples per segment at lower frequencies (i.e., *N =* 16 for 80, 250, and 500 Hz) and more samples per segment at higher frequencies (i.e., *N =* 32, 64, 128, and 256 for 1, 2, 4, and 8 kHz).

#### 2.2.3. Spectral Analysis

The steady-state frequency responses of the ear canal pressure, as well as the umbo and SFP displacements, were quantified using a fast Fourier transform (FFT) of the simulated time-domain signals:

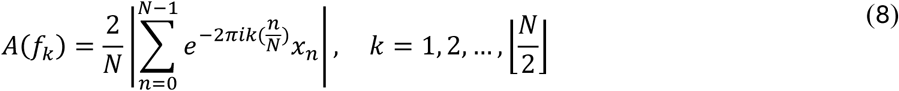

where *x_n_* represents the discrete time-domain signal, *n* is the sample index, *N* is the total number of samples, *k* is the frequency bin index associated with the discrete frequency *f_k_*, and 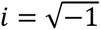. To eliminate transient effects, only the second half of each signal was retained for analysis, as suggested by the results of the steady-state determination (refer to Section 3.4). The frequency vector was defined as *f_k_* = *kF_s_/N*, where *F_s_* is the sampling frequency. For each case, the response amplitude at the excitation frequency, *f_exc_*, was extracted by identifying the closest frequency bin, denoted as A(*f_exc_*).

To quantify nonlinear and broadband spectral content, the total spectral distortion (TSD) was computed as:

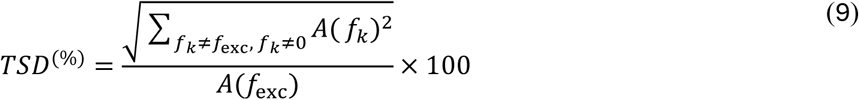

where *f_exc_* corresponds to the excitation frequency, and the DC component (*k* = 0), and both were excluded. This metric captures the total energy distributed across all frequency components other than the fundamental (excitation) frequency, including harmonics (integer multiples of the fundamental frequency), subharmonics (fractional multiples), and other nonlinear modulated components, normalized by the fundamental response. The analysis was applied to the ear canal pressure, umbo displacement, and SFP displacement signals to evaluate both the linear response amplitude and deviations from a purely sinusoidal pattern.

For both A(*f_exc_*) and TSD measures in the umbo and SFP displacement signals, the contributions from all three displacement components in the LM, AP, and SI directions were combined using the vector magnitude:

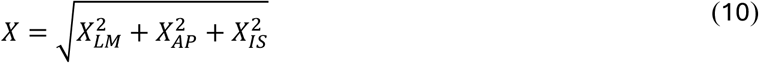

where *X* represents either A(*f_exc_*) or TSD, and *X_LM_*, *X_AP_*, and *X_IS_* denote the corresponding quantities computed from each directional component.

#### 2.2.4. Incremental Nonlinearity Index Analysis

Nonlinear response behavior was characterized by analyzing the displacement magnitude versus pressure relationship in logarithmic space, assuming *u ∝ p^γ^*. Therefore, the nonlinearity index, *γ*, was defined as the logarithmic slope of displacement with respect to ear canal pressure, computed as:

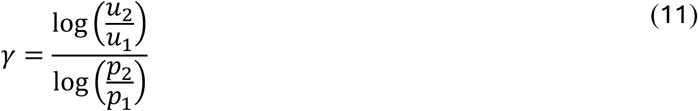

where *p_1_* and *p_2_* represent ear canal pressure at 1-mm distance from the umbo at a reference level and the level of interest, respectively. Similarly, *u_1_* and *u_2_* are their corresponding displacement magnitudes of either the umbo or SFP. A value of *γ* = 1 indicates linear behavior, while *γ* < 1 and *γ* > 1 correspond to compressive and expansive nonlinear responses, respectively.

An incremental nonlinearity formulation was used, where the index was computed between two consecutive stimulus levels, providing a local estimate of the input–output slope and capturing variations in response over fixed input increments (i.e., 10 dB SPL). For visualization, the nonlinearity index was expressed on a decibel scale as γ_dB_ = 20log_10_(γ), such that a linear response is centered at 0 dB. A threshold of ±0.5 dB was used to distinguish expansive and compressive nonlinearities from linear behavior.

## 3. Results

In this section, we present the transient responses and full-field displacements of the middle ear model at representative low-, mid-, and high-frequency conditions (0.5, 4, and 8 kHz, respectively) for two different levels corresponding to lower and higher sound intensities. Subsequently, we present the steady-state responses for all the frequencies and levels investigated.

### 3.1. Ear Canal Pressure

Figure 2 shows the time-domain ear canal pressure (0–16 ms), measured at 1 mm from the umbo, and the corresponding frequency-domain representation obtained from the second half of the signal (i.e., 8–16 ms). Panels (a), (b), and (c) present the results at input frequencies of 0.5, 4, and 8 kHz, respectively. The results are shown for a lower level of 120 dB SPL for input frequencies of 0.5, 4, and 8 kHz, and for higher levels of 170 dB SPL at 0.5 kHz and 150 dB SPL at 4 and 8 kHz. At 120 dB SPL, the amplitudes of the fundamental peak in the frequency domain were 20.1 Pa (120.0 dB SPL), 21.6 Pa (120.7 dB SPL), and 26.8 Pa (122.5 dB SPL) for 0.5, 4, and 8 kHz, respectively. At 150 dB SPL, the amplitudes were 635 Pa, corresponding to 150.0 dB SPL (not shown in Fig. 2), 634 Pa (150.0 dB SPL), and 1005 Pa (154.0 dB SPL) for 0.5, 4, and 8 kHz, respectively. Moreover, at 170 dB SPL and 0.5 kHz, the pressure amplitude was 6342 Pa (170.0 dB SPL). Distortion was noticeable at low and high levels and for 4 and 8 kHz, but not for 0.5 kHz. Figure 2 shows that the measured pressures were close to the prescribed pressure, and the differences are due to reflections from the TM.

**Fig. 2.**
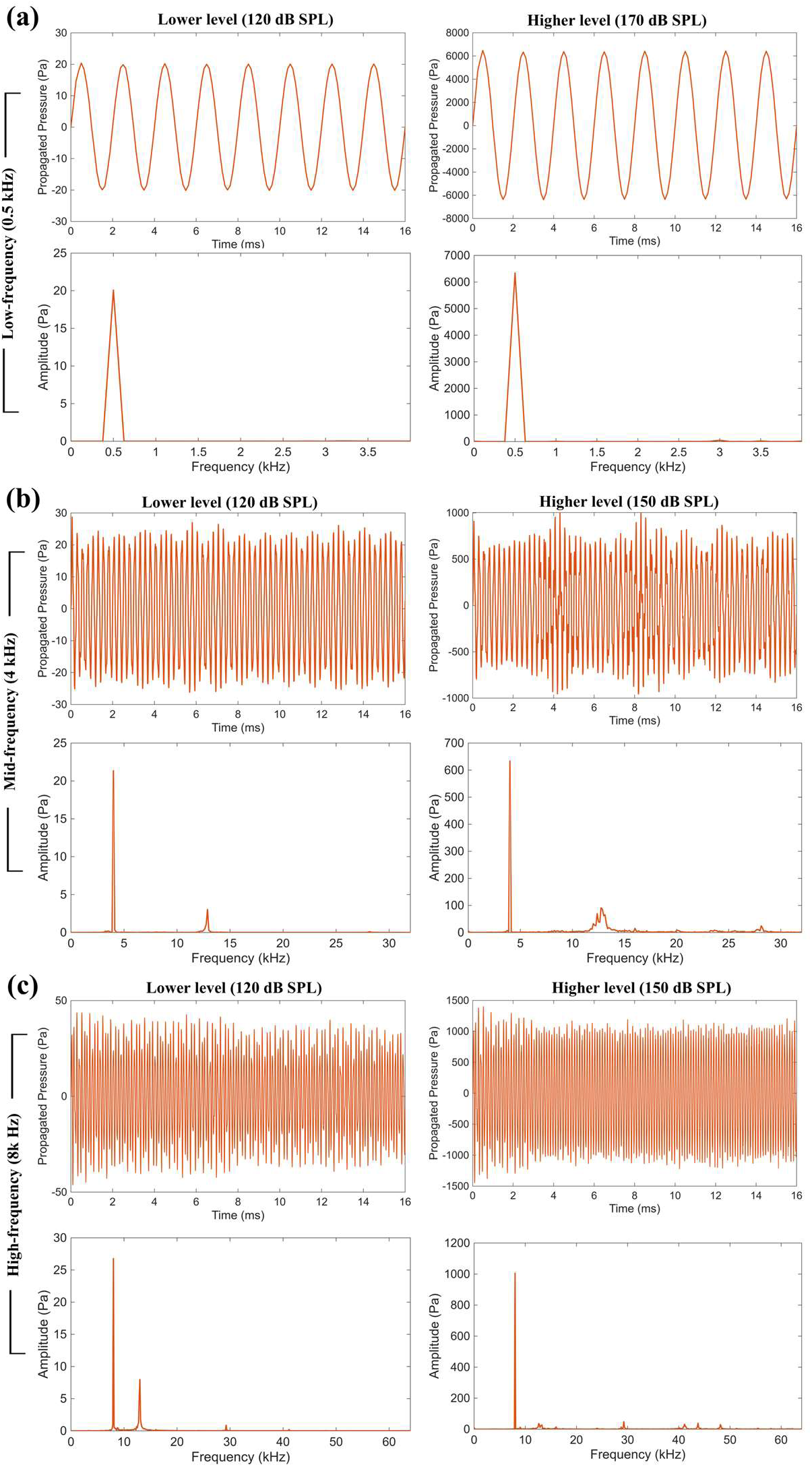
Ear canal pressure measured 1 mm from the umbo at representative low-, mid-, and high-frequency conditions (i.e., 0.5, 4, and 8 kHz, respectively) across rows in the time domain (0–16 ms), alongside the corresponding FFT plots obtained from the second half of the signal (8–16 ms). Results are presented at a lower sound pressure level (120 dB SPL) in the left column and a higher sound pressure level (170 dB SPL for 0.5 kHz and 150 dB SPL for 4 and 8 kHz) in the right column. The time-domain plots show a nearly perfect pure-tone pressure response at low frequency and slight waveform distortions at mid and high frequencies. The corresponding frequency-domain plots at mid and high frequencies also show spectral distortion products in addition to the fundamental peaks at 4 and 8 kHz. Note the different frequency scales in the lower rows for each frequency plot. The reported pressure is the total pressure consisting of the sum of incident and reflected pressures

### 3.2. Three-Dimensional TM and Ossicle Displacements

Figure 3 shows the three-dimensional full-field displacement magnitudes of the middle-ear structures and transient motion of the umbo and SFP for representative frequencies and levels. Figure 3a demonstrates the displacement distribution across the surface of the TM at a peak instance of the excitation pressure during the last cycle of the simulation (i.e., at t = 15.5 ms). At 0.5 kHz, the maximum displacement occurred in the posterior–superior (PS) quadrant of the TM, located between the umbo and the annulus. The distribution of displacement magnitudes was relatively smooth and continuous, indicating a simple mode shape, with the amplitude gradually decreasing toward the annulus due to the fixed boundary constraint and toward the umbo due to the mechanical coupling with the ossicular chain. The maximum local displacement magnitude increased from 2.00 μm to 395.4 μm as the level increased from 120 to 170 dB SPL. At 4 kHz, the TM exhibited more complex mode shapes, with maximum local magnitudes of 0.89 μm and increasing to 125.7 μm when the level increased from 120 to 150 dB SPL. Similarly, at 8 kHz, a higher-order mode shape was observed, with maximum magnitudes of 3.40 μm and 39.38 μm at 120 dB SPL and 150 dB SPL, respectively.

**Fig. 3.**
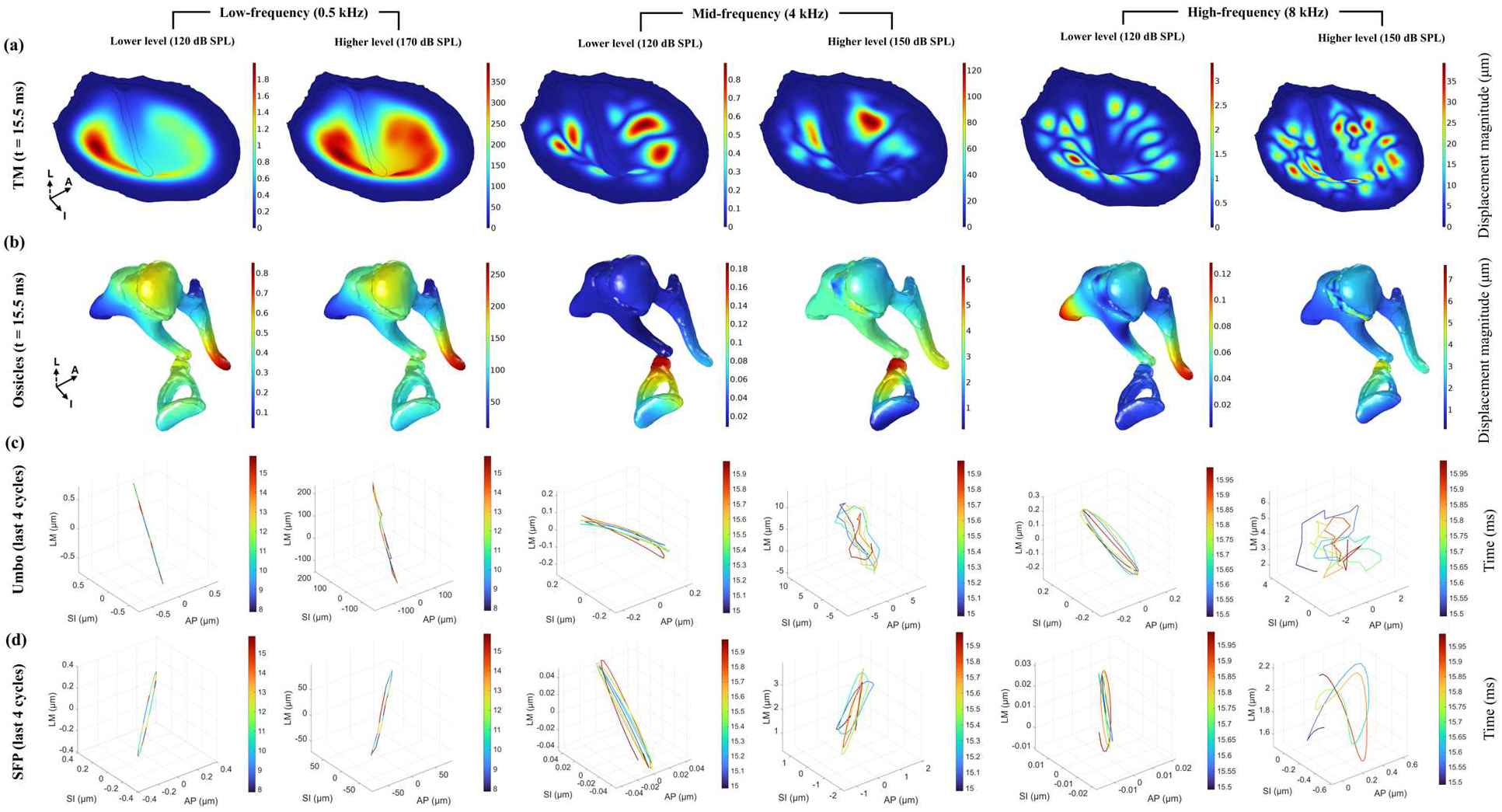
Displacement propagation through the TM and ossicular chain in the middle ear at representative low- (0.5 kHz), mid- (4 kHz), and high-frequency (8 kHz) conditions for lower- and higher-level pure-tone pressure excitations. Subplots in rows (a) and (b) demonstrate the full-field displacement magnitude at a peak-pressure instance during the final cycle of the simulation. At low frequency, the TM exhibits a low-order mode shape with predominantly global piston-like motion across the ossicular chain, while higher-order modes with rocking motions are visible at higher frequencies. Animations of the full-field displacement of the TM and ossicular chain for these representative conditions are provided in SM. Subplots in rows (c) and (d) show the umbo and SFP displacement components in the LM, SI, and AP directions during the last four cycles of the simulation. At low frequency, the displacement is dominated by the LM direction (i.e., piston-like motion), whereas at higher frequencies, the contributions of SI and AP displacements become more pronounced. Note the different scale bars for each subplot. Animations of the 3D motions of the umbo and SFP under the representative conditions are also provided in SM

Figure 3b presents the displacement magnitude fields across the ossicular chain (from the umbo to the SFP) for the same frequencies and levels at the same time point (i.e., t = 15.5 ms). At 0.5 kHz, the displacement field was relatively uniform across the stapes, consistent with its near-rigid-body motion and stiffness-dominated behavior at low frequencies. In contrast, at 4 kHz and 8 kHz, larger spatial variations in displacement magnitude, particularly across the stapes, were observed, suggesting increased relative motion across the ossicular joints as inertial effects became more pronounced. Bending associated with relative ossicular motion was most evident near the ISJ at 4 kHz and the IMJ at 8 kHz.

The magnitude range across the ossicular chain was generally lower than that observed for the TM. At 0.5 kHz, the displacement magnitude reached 0.85 μm at 120 dB SPL and 268.29 μm at 170 dB SPL, with the maximum magnitude occurring at the umbo and gradually decreasing along the ossicular chain. The displacement magnitude on the SFP reached up to around 0.5 μm and 150 μm for 120 dB SPL and 170 dB SPL, respectively. At 4 kHz, a different motion pattern was observed due to a phase shift, resulting in larger displacements near the ISJ, with maximum magnitudes of 0.19 μm at 120 dB SPL and 6.55 μm at 150 dB SPL. At 8 kHz, the displacement modes became more complex, exhibiting frequency- and level-dependent behavior. Maximum displacement magnitudes of 0.13 μm occurred around the umbo and PIL at 120 dB SPL, while the magnitudes increased up to 7.66 μm around the joints, particularly near the IMJ, at 150 dB SPL.

The rows in Fig. 3c and 3d present the three-dimensional motion of the umbo and SFP, along the LM, SI, and AP directions, during the last four cycles of the time-domain simulations. At low frequency (0.5 kHz), the motion of both the umbo and SFP was primarily along the LM direction at both stimulus levels, exhibiting an approximately piston-like response with motion largely confined along that direction. At higher frequencies (4 and 8 kHz), increased contributions from the SI and AP directions were observed, indicating a transition toward multi-directional motion. At 120 dB SPL, the trajectories remained largely planar, suggesting predominantly two-dimensional motion. However, at higher stimulus levels (150 dB SPL), the trajectories became more complex and exhibited fully three-dimensional motion, reflecting the contribution of multiple vibration modes and increased modal coupling.

It should be noted that the subplots at 4 and 8 kHz do not represent the maximum displacement instances; instead, they correspond to the time instances at which the maximum ear canal pressure occurred. Animations of the full-field motion patterns, as well as the nodal displacements of the umbo and SFP, are provided in the Supplementary Materials (SM) for a more complete representation of the full-field transient responses.

### 3.3. Time-and-Frequency Domain Motion Analysis

Figure 4 shows the time-and-frequency domain umbo displacement components in the AP, SI, and LM directions over a 16-ms period, with the LM direction highlighted in blue solid lines. The LM direction corresponds to the piston-like direction and represents the dominant component of motion in most cases, particularly for frequencies up to 2 kHz. Panels (a), (b), and (c) present the results at three representative input frequencies of 0.5, 4, and 8 kHz, respectively, each at two different stimulus levels. The top rows show the time-domain results, and the bottom rows show the corresponding FFT-derived frequency-domain results, obtained from the second half of the time-domain responses (i.e., 8–16 ms).

**Fig. 4.**
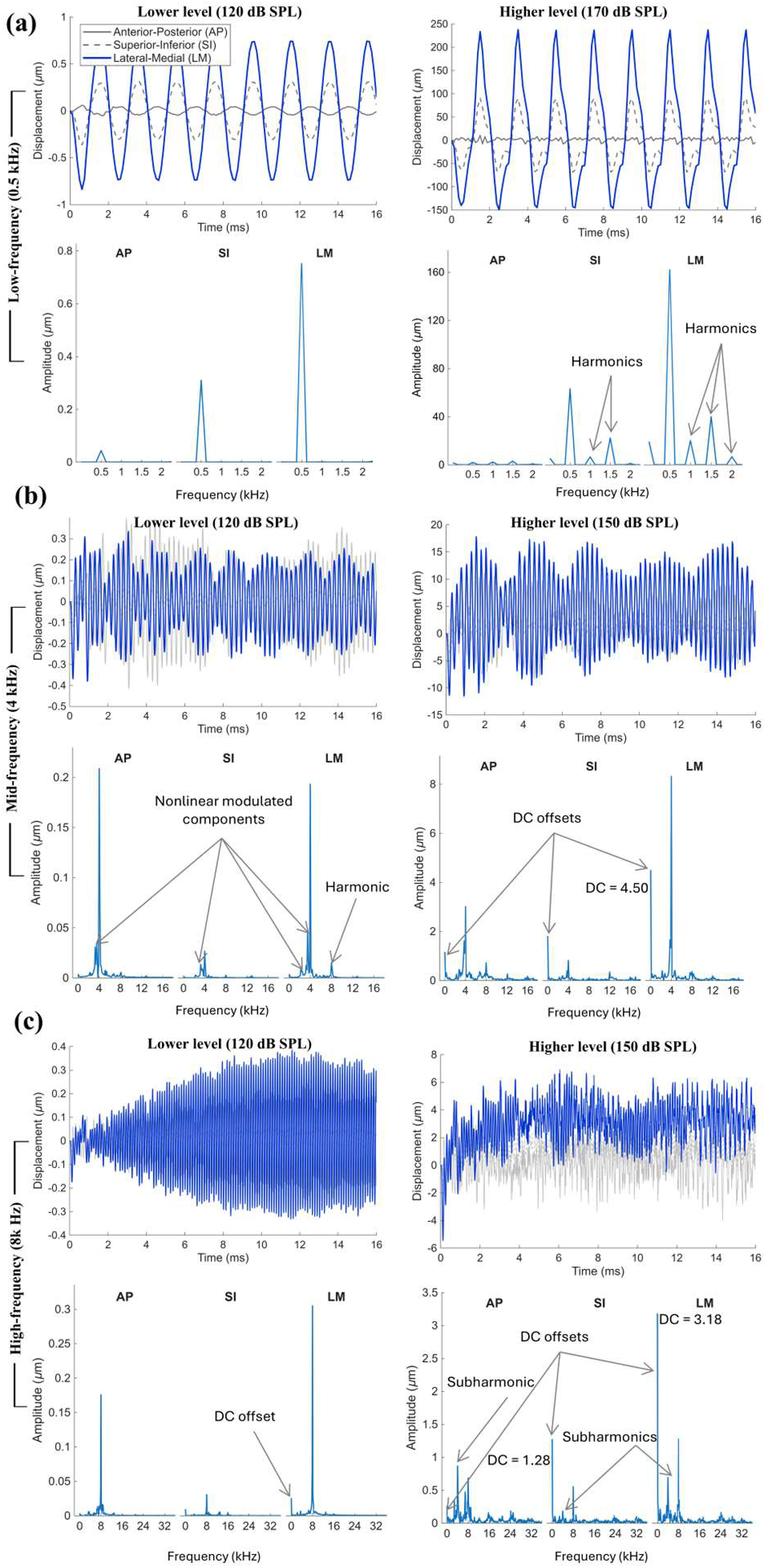
Umbo displacement at representative (a) low- (0.5 kHz), (b) mid- (4 kHz), and (c) high-frequency (8 kHz) conditions for lower- and higher-level pure-tone pressure excitations, shown in the time domain over 16 ms (top row) alongside the corresponding FFT plots obtained from the second half of the signal (8–16 ms) (bottom row). The piston-like LM component is highlighted in blue, while the SI and AP components are shown in gray for overall comparison in the time-domain plot; three motion components are clustered in the FFT-derived frequency-domain plot as labeled. At low-frequency, lower-level excitation, the displacement response is a pure-tone sinusoidal waveform. At low frequency and higher-level excitation, harmonic components appear in the response, although the motion remains periodic. At mid frequency, the time-domain response exhibits amplitude modulation characterized by a beating envelope, arising from the interaction of multiple nonlinear modulated frequency components visible in the FFT plots. At mid frequency and high level, DC offsets start to emerge at all three motion components. At high frequency and lower-level excitation, the response shows a transient build-up toward a nearly pure-tone sinusoidal waveform in the time-domain plot, and a small DC offset in the LM direction is revealed in the FFT plot. At high frequency and higher-level excitation, the response exhibits pronounced DC offsets (primarily in the LM direction) together with amplitude modulation associated with an irregular envelope. In the corresponding FFT plots, these DC offsets are clearly seen as peaks at f = 0, while the amplitude modulation is associated with the generation of subharmonics and modulated components

At low frequency (0.5 kHz) and at a low level of 120 dB SPL, the umbo displacement in the time domain exhibited sinusoidal motion patterns in all directions, with the LM direction representing the dominant motion component. The frequency-domain results indicate an amplitude of 0.75 μm in the LM direction, compared to 0.31 μm in the SI direction and only 0.04 μm in the AP direction. At the same frequency but at the highest investigated level (170 dB SPL), the motion in the AP direction became negligible, while the SI motion remained minor compared to the LM motion (63.25 μm versus 161.89 μm). In addition, several harmonic components appeared in the response, with peak amplitudes of 20.14, 40.03, and 6.78 μm (for the LM component) occurring at 1, 1.5, and 2 kHz, respectively. The third harmonic at 1.5 kHz was considerably large (24.7% relative to the fundamental LM response), with the second harmonic at 1 kHz being the second largest (12.4%), indicating that the distortion was predominantly odd-order, with a weaker even-order component.

At mid frequency (4 kHz), the responses exhibited amplitude modulation with beating-envelope patterns in the time domain. In the frequency domain, these signals were associated with multiple modulated components. These components corresponded to the fundamental peak at the excitation frequency and additional FFT peaks that occurred at frequencies offset from the excitation frequency, rather than at harmonic or subharmonic frequencies. At the lower level (120 dB SPL), the AP component was more substantial and comparable to the LM component (0.21 μm versus 0.19 μm), while the SI component remained almost negligible (0.03 μm), indicating an approximately planar motion within the LM–AP plane. In contrast, at the higher level (150 dB SPL), the LM component became the dominant motion component, with an amplitude of 8.32 μm compared to 3.02 μm and 0.82 μm in the AP and SI directions, respectively, and with a predominant LM piston-like pattern similar to the low-frequency condition. Although the modulated components were not substantial compared to the fundamental peak, they contributed to the overall displacement response and appeared as envelope effects in the time-domain signals. Note that now we see DC offsets start to emerge. As discussed next, these DC offsets become more significant at higher frequencies.

At high frequency (8 kHz) and low level (120 dB SPL), the time-domain responses exhibited more complex patterns, the most striking of which is the transient build-up in the first 6-8 ms for a 120 dB SPL input tone. At this level, the dominant motion components were LM (0.30 μm) and AP (0.18 μm), while the SI component remained almost negligible (0.03 μm), indicating a nearly planar motion within the LM–AP plane, similar to the response observed at the same level for the mid-frequency case.

At high frequency (8 kHz) and high level (150 dB SPL), the transient build-up was approximately 2 ms and thus appears faster than at the low level. In addition, a fully three-dimensional motion was observed, with amplitudes of 1.28, 0.69, and 0.55 μm in the LM, AP, and SI directions, respectively. Moreover, additional distortions emerged in the displacement responses in the frequency domain. The DC offsets, seen in the time-domain traces, are now quite large, particularly in the LM direction. The DC offsets were 3.18, 0.27, and 1.28 μm in the LM, AP, and SI directions, respectively. The DC offsets in the LM and SI directions were even greater than the corresponding amplitudes at the excitation frequency. Additonally, subharmonics appeared, with amplitudes of 0.69 μm in the LM direction, 0.87 μm in the AP direction, and 0.18 μm in the SI direction. The subharmonic peak in the AP direction was slightly larger than the corresponding excitation-frequency amplitude. The absence of harmonics and the emergence of subharmonics and large DC offsets suggest a shift in the underlying dynamics at the umbo for the 8 kHz high frequency and 150 dB SPL high level input. This phenomenon is not observed at the lower levels and frequencies thus far.

Similar to Fig. 4 for the umbo, Fig. 5 shows the time-domain SFP displacement at 0.5, 4, and 8 kHz in panels (a), (b), and (c), respectively. As in Fig. 4, the frequency-domain responses (below the time-domain responses) represent only the second half of each signal (i.e., 8–16 ms).

**Fig. 5.**
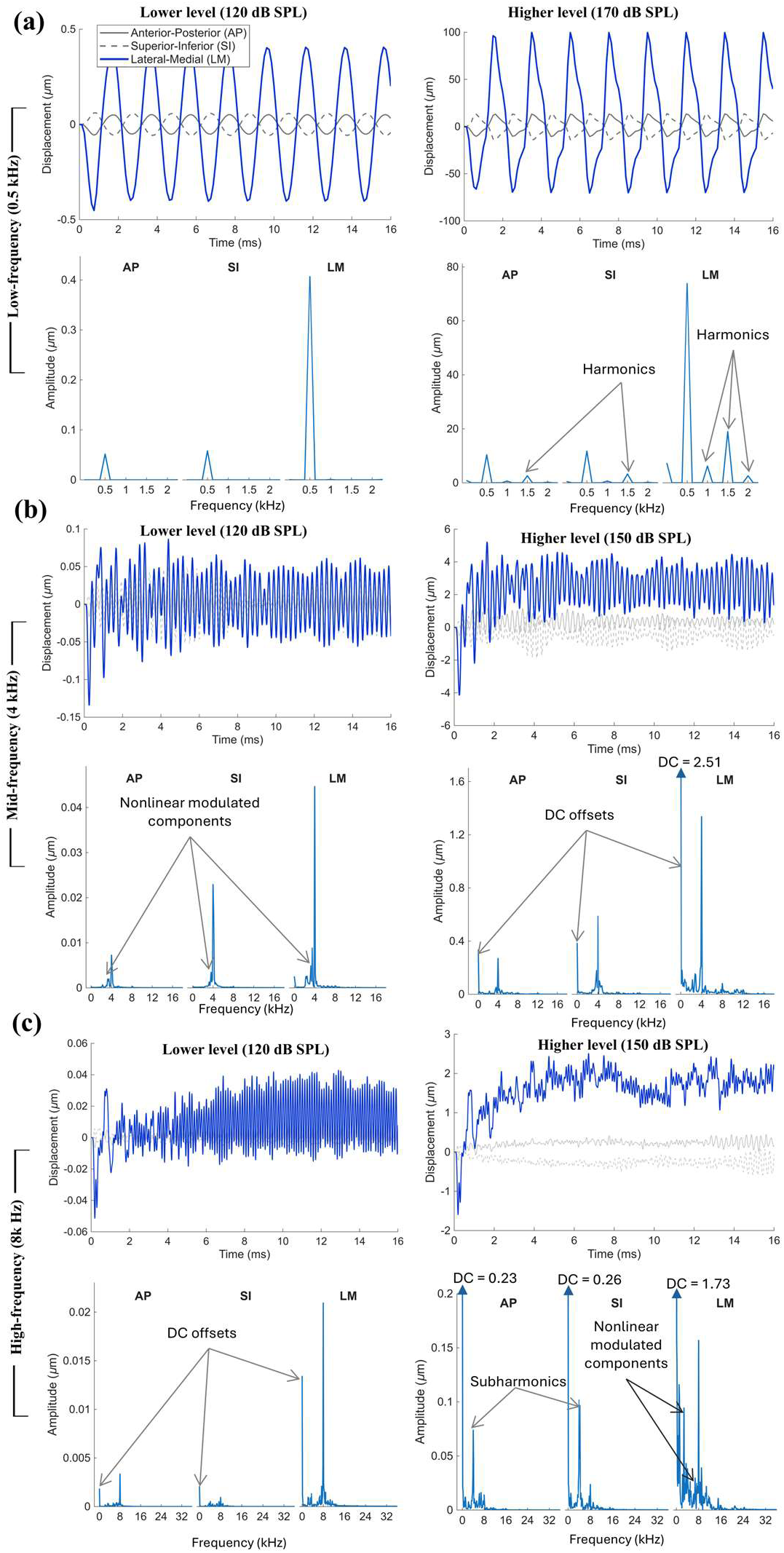
Same format as Fig. 4, but for SFP displacement. At low-frequency, lower-level excitation, the response exhibits a pure-tone sinusoidal waveform, whereas at low-frequency, higher-level excitation, harmonic components appear. At mid-frequency conditions, amplitude modulation is generated in the response, appearing as beating envelopes in the time-domain plots and nonlinear modulated components in the FFT plots. At high-frequency, lower-level excitation, the response exhibits a transient build-up toward a nearly pure-tone sinusoidal waveform with a small DC shift. At higher levels for both mid- and high-frequency excitations, substantial DC shifts are produced, predominantly visible in the LM direction

At low frequency (0.5 kHz) and low-level stimulation (120 dB SPL), the SFP displacements in all directions exhibited perfect pure-tone sinusoidal behavior, with amplitudes of 0.41, 0.05, and 0.06 μm in the LM, AP, and SI directions, respectively. This suggests a predominantly piston-like motion due to the negligible AP and SI components compared to the LM component. At 170 dB SPL, the LM motion remained the dominant component (73.84 μm) compared to the AP (10.38 μm) and SI (11.76 μm) components. Harmonic components appeared in all directions. For example, harmonics with amplitudes of 6.21, 19.00, and 2.60 μm appeared at 1, 1.5, and 2 kHz, respectively, in the LM direction. Although these harmonics were relatively small compared to the SFP displacement amplitude at the excitation frequency, they partially contributed to the time-domain responses and appeared as distortions deviating the signal from purely sinusoidal shapes. Nevertheless, the SFP displacement remained periodic, even at the highest level, i.e., 170 dB SPL.

At mid-frequency (4 kHz) and lower-level stimulation (120 dB SPL), envelope modulation patterns appeared in the time-domain responses, associated with nonlinear modulated components in the frequency domain. The fundamental amplitudes in the frequency domain were 0.045, 0.023, and 0.007 μm in the LM, SI, and AP directions, respectively. This pattern indicated an approximately LM–SI planar motion, in contrast to the umbo response, where an LM–AP planar motion was observed. At the higher level (150 dB SPL), the fundamental amplitudes increased to 1.336 μm in the LM direction, to 0.587 μm in the SI direction, and to 0.269 μm in the AP direction, indicating a response closer to a predominantly piston-like motion. In addition, the LM response exhibited a DC offset greater than the fundamental amplitude (2.510 μm versus 1.336 μm), with additional modulated components appearing near the DC component and on the lower-frequency side of the excitation frequency.

At high frequency (8 kHz), the lower-level stimulation (120 dB SPL) resulted in an almost piston-like motion, with a fundamental amplitude of 0.021 μm in the LM direction and negligible components in the AP (0.003 μm) and SI (almost zero) directions. At the higher-level stimulation (150 dB SPL), a more complex three-dimensional motion pattern was observed. The displacement in the LM direction had a fundamental amplitude of 0.157 μm, accompanied by substantial nonlinear modulated components, particularly on the lower-frequency side of the excitation frequency. In addition, subharmonics with amplitudes of 0.102 μm in the SI direction and 0.074 μm in the AP direction were observed. The subharmonic amplitudes in the AP and SI directions were greater than the corresponding fundamental amplitudes. These distortions appeared as modulation patterns in the time-domain responses, causing the SFP to move at frequencies substantially lower than the excitation frequency. Moreover, the responses exhibited substantial DC offsets (1.732 μm in LM, 0.269 μm in SI, and 0.234 μm in AP directions), all of which were greater than the corresponding fundamental amplitudes and the observed subharmonic amplitudes. The LM DC offsets are generally positive such that the SFP is offset in the lateral direction away from cochlear fluids.

### 3.4. Steady-State Determination

Figure 6 presents delay-embedded Poincaré maps of umbo (top row) and SFP (bottom row) motion in the LM direction at 0.5, 4, and 8 kHz in panels (a), (b), and (c), respectively, at two sound pressure levels: a lower level (left) and a higher level (right). The color scale represents the temporal progression over the final cycles of the steady-state response. For the Poincaré plots, each excitation cycle was sampled at 16 equally spaced time points, defined as *t*_m,j_ = (*m* + *j*/16)/*f*_exc_ where *m* denotes the cycle number and *j* = 0,…, 15 denotes the sample within each cycle. Thus, the plots included 128, 1024, and 2048 samples corresponding to 8, 64, and 128 cycles at 0.5, 4, and 8 kHz, respectively. Asterisks and squares denote the beginning and end of each trajectory, respectively, while arrows indicate the direction of time progression. At low frequency (0.5 kHz), both umbo and SFP responses exhibited nearly elliptical trajectories with distinct point clusters at 120 dB SPL, consistent with predominantly harmonic motion (Fig. 6a). At the higher excitation level (170 dB SPL), the trajectories expanded significantly in amplitude (see values on the x- and y-axes) and became non-elliptical, indicating distortions due to the presence of higher-order harmonics arising from model nonlinearities. Nevertheless, at both levels, the responses remained highly periodic (except during the first cycle, represented by dark-blue points), with samples from corresponding time instances in successive cycles repeatedly occupying the same coordinates and forming 16 distinct clusters corresponding to the 16 sampling points per cycle. As an example, these 16 clusters are indicated by dotted orange circles for the umbo response at 120 dB SPL. This observation indicates that the middle-ear response reached steady state for low-frequency input after the first cycle.

**Fig. 6.**
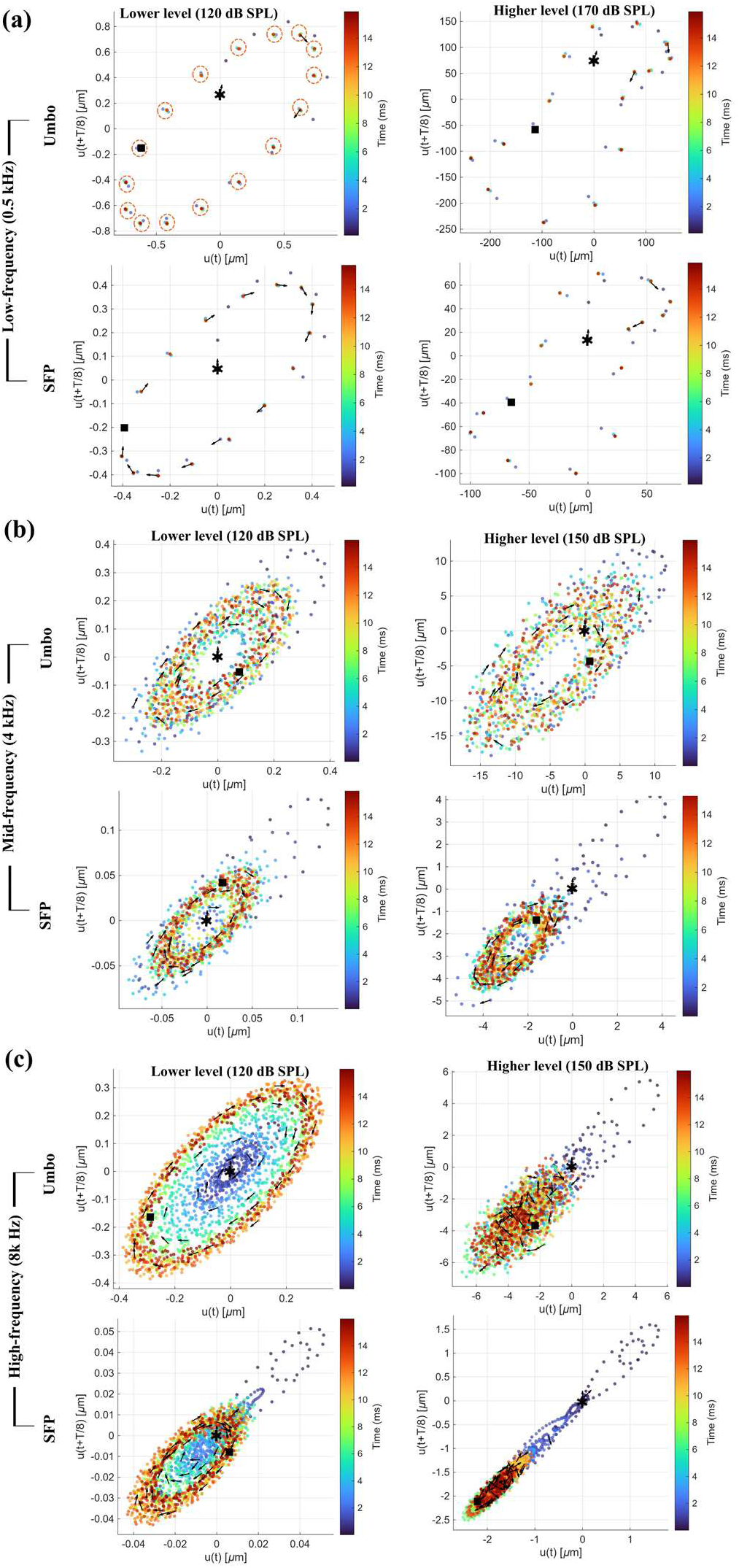
Poincaré maps of LM displacements for both the umbo (top) and SFP (bottom) at representative (a) low- (0.5 kHz), (b) mid- (4 kHz), and (c) high-frequency (8 kHz) conditions, presented at a lower sound pressure level (120 dB SPL) and a higher sound pressure level (170 dB SPL for 0.5 kHz and 150 dB SPL for 4 and 8 kHz). At low-frequency excitations, the Poincaré maps form well-defined closed-loop orbits from the beginning to the end of the response (blue-to-red regions), indicating a periodic steady-state response. The lower-level excitation exhibits a thin, nearly elliptical trajectory representing a predominantly single-frequency response, whereas the higher-level excitation becomes more distorted due to harmonic generation. At higher frequencies, the Poincaré maps initially exhibit elongated transient structures (blue regions) that gradually evolve toward more established closed-loop elliptical patterns at later times (red regions), indicating convergence toward a steady-state response within the second half of the signals (8–16 ms). The plot is at time *t* along the horizontal axis versus at *t* = *t* + *T/8* on the vertical axis

At higher frequencies (4 and 8 kHz), the Poincaré maps for both the umbo and SFP exhibited more irregular and dispersed patterns, in which the distinct 16-cluster structure was no longer clearly distinguishable. In most cases, the samples were initially distributed in elongated and sporadic patterns (blue regions in Fig. 6b and 6c) and gradually evolved into more structured elliptical trajectories toward the middle and end of the response (green-to-red regions), indicating convergence toward a steady-state pattern. At 8 kHz and 120 dB SPL, the elliptical structure expanded over time, with later samples (red regions) exhibiting larger amplitudes than earlier samples (blue regions). In addition, the centers of the Poincaré maps at high-frequency, high-level stimulations shifted noticeably, indicating the presence of a DC offset, particularly visible at 4 kHz and 150 dB SPL & 8 kHz and 150 dB SPL. These Poincaré maps demonstrate that, despite the complex and irregular appearance of the time-domain responses, the later portions of the high-level, high-frequency responses converge toward stable dynamical orbits.

Yet another way to look at the convergence of the response is to look at the normalized Euclidean distance (difference) between adjacent signal segments (Equation 7). This is shown in Fig. 7 for both umbo (top row) and SFP (bottom row) displacement responses in the LM direction at different frequencies (80 Hz to 8 kHz) and sound pressure levels of 120, 150, and 170 dB SPL in Figs. 7a, 7b, and 7c, respectively. The horizontal axis indicates the segment number relative to the preceding segment, starting from segment 2. At the highest level (170 dB SPL), only frequencies up to 500 Hz were included. In all cases, the signals were divided into eight segments, except for the low-frequency cases (i.e., 80 and 250 Hz), where only four segments were available. For both umbo and SFP displacements, the normalized differences between adjacent segments generally increased with frequency, indicating greater temporal variability at higher frequencies. However, the responses at 2 kHz exhibited the largest segment differences, exceeding those at 4 and 8 kHz in most cases.

**Fig. 7.**
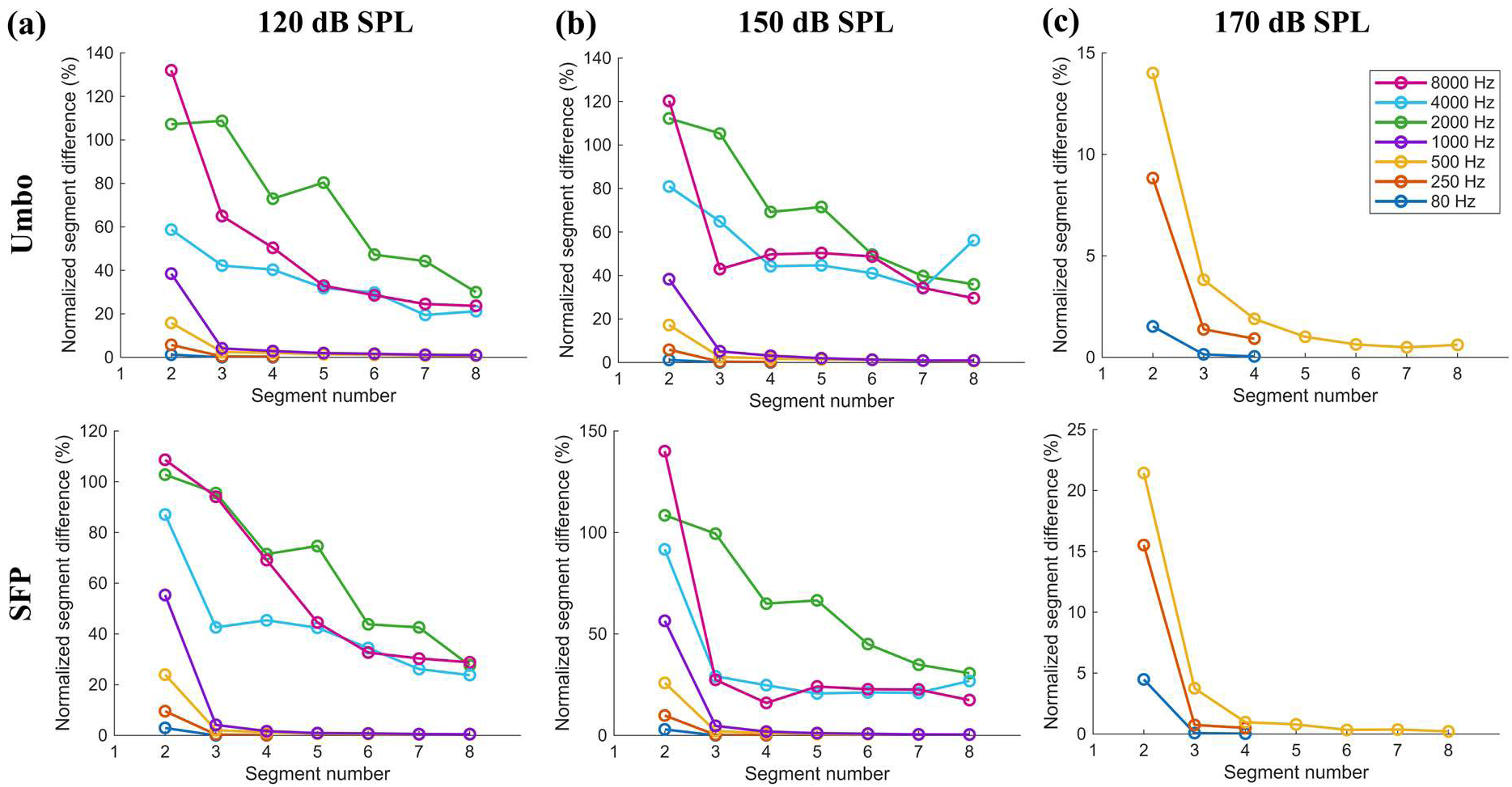
Normalized segment differences of time-domain LM displacements for both the umbo (top) and SFP (bottom) at frequencies ranging from 80 Hz to 8 kHz and three representative sound pressure levels: (a) 120, (b) 150, and (c) 170 dB SPL. The time-domain responses were divided into eight equal-length consecutive segments, except for the 80 Hz and 250 Hz cases, where only four equal-length segments were used. For each pair of adjacent segments, a normalized Euclidean distance was computed between the corresponding signal vectors. In all frequency–level cases, a convergence plateau is observed starting from segment 3 or 4, indicating a steady-state response during the subsequent cycles of the simulation. Note that at the highest level (170 dB SPL), only three frequencies up to 500 Hz were shown

At high frequencies (2–8 kHz), the Euclidean distances started from values around or above 100% for the first comparison (segment 2 versus segment 1), then decreased to below 40% in the subsequent segments, typically reaching a plateau from segment 3 onward. At low-frequency conditions, the Euclidean distances initially were about 2% at 80 Hz, 5% at 250 Hz, 15% at 500 Hz, and 40% at 1 kHz, and converged to values below 5% in all cases, generally from segment 3 or 4 onward.

Overall, the normalized Euclidean distances decreased and then plateaued in the later segments (typically after the first quarter of the signal) for all cases, indicating that the responses had reached a steady state. For low-frequency cases (up to 1 kHz, below the resonance frequency of the middle ear), the responses converged to a periodic steady state (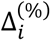 ≈ 0). In contrast, at frequencies of 2 kHz and above (above the resonance frequency), the normalized distances plateaued despite the responses not converging to a perfectly periodic cycle, indicating a non-periodic steady state.

### 3.5. Steady-State Responses

Considering the steady-state determination results, only the second half of the ear canal pressure and the umbo and SFP displacement signals (8–16 ms) were used as representative of the model steady-state behavior. Figure 8 plots the steady-state responses of the model obtained from the fundamental peaks of the spectral responses at the investigated frequencies and stimulus levels. Figure 8a shows the pressure amplitude (y-axis) measured near the TM, 1 mm from the umbo, with respect to the input pressure (x-axis), suggesting a linear increase (parallel to the linear growth line) of the ear canal pressure near the TM with the input pressure. The difference between the ear canal pressure near the TM and the input pressure (Fig. 8-a right column) was less than 8% across most frequencies. The exception was at 8 kHz, where the pressure amplitude increased by approximately 35% at 110 and 120 dB SPL and by approximately 60% at 130, 140, and 150 dB SPL.

**Fig. 8.**
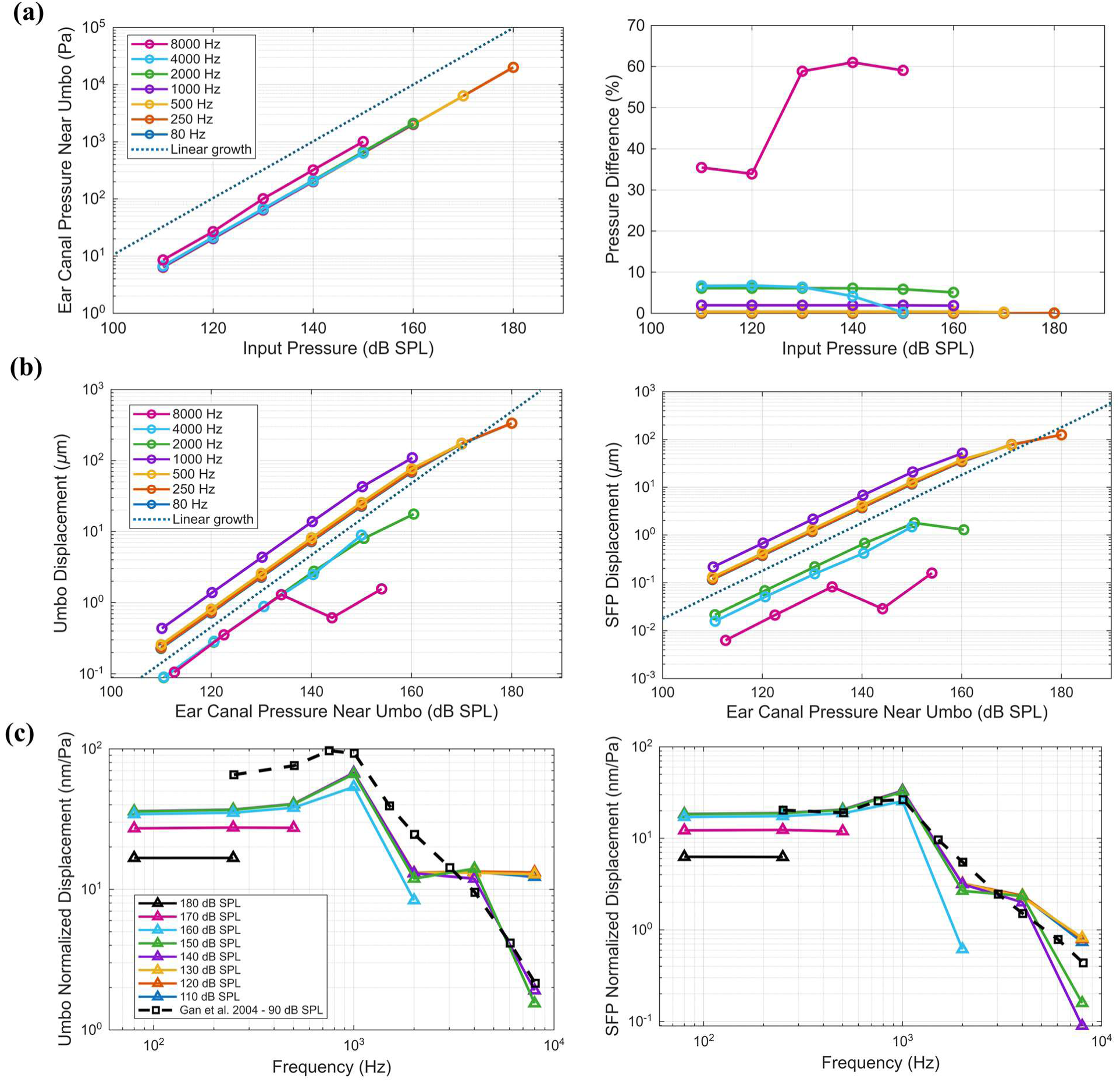
Steady-state responses of the model computed from the FFT of the second half of the time-domain signals for all investigated frequencies and sound pressure levels: (a) ear canal pressure measured at 1 mm from the umbo versus the prescribed input SPL, compared with a linear growth indicator (left). Alongside, in the right column, its difference from the input SPL; (b) umbo (left) and SFP (right) displacement magnitudes versus the input SPL for different frequencies, compared with linear growth indicators; and (c) normalized displacement magnitudes of the umbo (left) and SFP (right) as a function of frequency for different SPLs, compared with the experimental data from *(Gan et al. 2004a)*, shown with black dashed lines

Figure 8b shows the umbo (left) and SFP (right) displacement magnitudes from three-dimensional components (Eq. 10) as a function of the ear canal pressure obtained from the model near the umbo. At low frequencies (80 Hz to 500 Hz), the displacement magnitudes increased almost linearly from 110 to 150 dB SPL, ranging from approximately 0.25 μm to 70 μm for the umbo and from approximately 0.12 μm to 35 μm for the SFP. Beyond this range, the slope decreased, indicating a compressive nonlinearity, with maximum magnitudes of 335 μm for the umbo and 125 μm for the SFP at 180 dB SPL. At 1 kHz, the displacement magnitudes increased from 0.44 μm at 110 dB SPL to 109 μm at 160 dB SPL for the umbo, and from 0.21 μm at 110 dB SPL to 52 μm at 160 dB SPL for the SFP. This increase, compared to the lower frequencies (<1 kHz), is likely related to the proximity of 1 kHz to the reported resonance range of the middle ear, resulting in a larger displacement response than the neighboring investigated frequencies.

At higher frequencies (>1 kHz), the displacement magnitudes were generally lower. At 2 and 4 kHz, increasing the pressure level from 110 to 150 dB SPL increased the umbo displacement magnitudes from approximately 0.09 μm to 8.0 μm, while the SFP displacement magnitudes increased from approximately 0.02 μm to 1.6 μm. A noticeable compressive nonlinearity emerged at 160 dB SPL, where the displacement magnitudes reached 17.6 μm for the umbo and 1.29 μm for the SFP. At 8 kHz, a more irregular nonlinear behavior was observed starting from 130 dB SPL. The displacement magnitudes ranged from 0.105 μm at 110 dB SPL to 1.55 μm at 150 dB SPL for the umbo, and from 0.006 μm at 110 dB SPL to 0.16 μm at 150 dB SPL for the SFP. More details regarding the quantified nonlinearities are presented in the following subsection.

Figure 8c shows the normalized displacement magnitudes as a function of frequency for different sound pressure levels. The experimental data from (Gan et al. 2004a) measured at 90 dB SPL are shown with dashed lines for comparison (their peak-to-peak amplitudes were divided by two to make them comparable with the present results obtained from the fundamental peaks in the frequency domain). Overall, the normalized displacement magnitudes remained relatively consistent for levels up to 150 dB SPL and frequencies up to 4 kHz, suggesting linear responses of the model within these levels. At frequencies between 80 and 500 Hz, the normalized displacement magnitudes were approximately 35 nm/Pa for the umbo and 18 nm/Pa for the SFP. At 1 kHz, these values increased to approximately 67 nm/Pa and 33 nm/Pa for the umbo and SFP, respectively, due to the resonance of the middle ear. At higher frequencies, the normalized magnitudes decreased substantially.

At levels above 150 dB SPL, the normalized displacement magnitudes generally decreased, suggesting compressive nonlinearity. For example, at 170 dB SPL, the displacement magnitudes were approximately 27 nm/Pa for the umbo and 12 nm/Pa for the SFP, while at 180 dB SPL, the magnitudes decreased to approximately 17 nm/Pa for the umbo and 6 nm/Pa for the SFP. These values remained relatively consistent across the investigated frequency range below resonance.

### 3.6. Transmission Along the Ossicular Chain

To further examine how sound-induced vibrations are transmitted through the middle-ear ossicular chain at different frequencies and sound stimulus levels in our model, we calculated the overall SFP-to-umbo displacement transfer ratio (Fig. 9a), as well as the incus-to-umbo and SFP-to-incus transfer ratios (Figs. 9b and 9c, respectively). Motion of the incus was obtained at its lenticular process (LP). The reciprocal (i.e., 0.7) of the anatomical lever ratio of 1.4, computed as the ratio of the malleus to incus lever arms (O’Connell-Rodwell et al. 2024), is also shown as a horizontal dashed line for reference. Figure 9 also includes the experimentally obtained transfer ratios reported by (Goode et al. 1994; O’Connell-Rodwell et al. 2024) for comparison. Overall, our model predictions are comparable with the experimental data. Note that in (O’Connell-Rodwell et al. 2024), the measurements were obtained at the stapes head rather than the SFP.

**Fig. 9.**
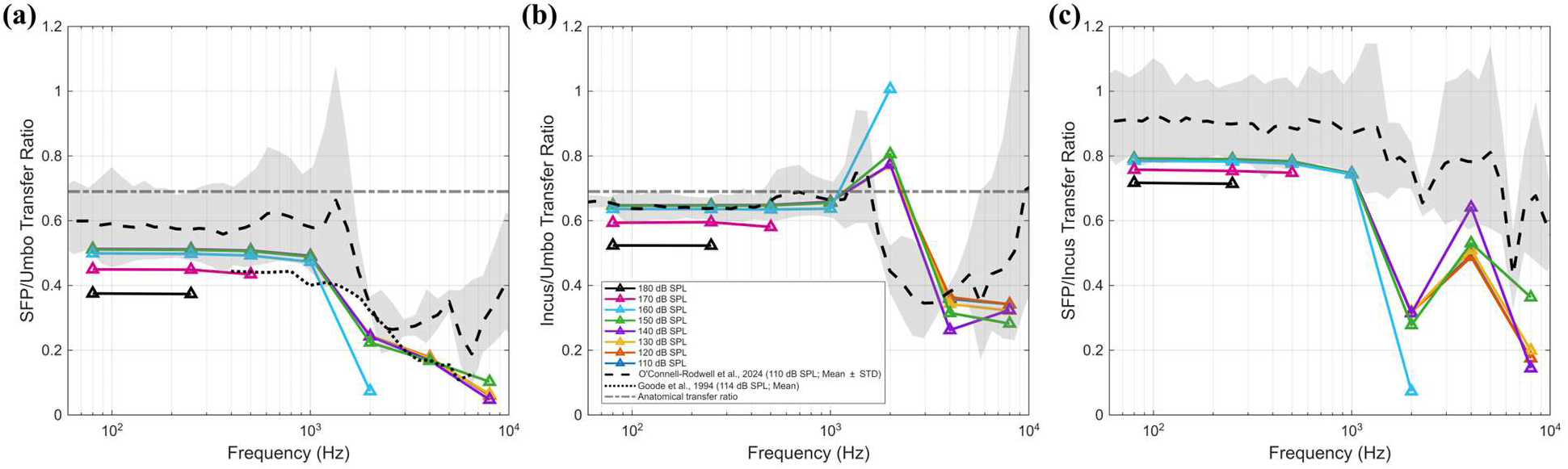
Overall SFP-to-umbo (a), incus-to-umbo component (b), and SFP-to-incus component (c) transfer ratios. The incus-to-umbo ratio corresponds to motion transmission through the ossicular lever arms and slippage across the IMJ, while the SFP-to-incus ratio corresponds to slippage across the ISJ and potential rotation and bending of the stapes. At low frequencies and lower sound pressure levels (generally up to 1 kHz and 160 dB SPL), both component transfer ratios remained nearly constant. At higher frequencies and SPLs, the transfer ratios generally decreased. Exceptions included a local peak in the incus-to-umbo ratio at 2 kHz and a local peak in the SFP-to-incus ratio at 4 kHz. The anatomical transfer ratio and experimental transfer ratios reported by *(Goode et al. 1994; O’Connell-Rodwell et al. 2024)* are shown for comparison

In Fig.9a, the overall SFP-to-umbo transfer ratio remained approximately constant at 0.51 up to 1 kHz and 160 dB SPL, and then decreased to about 0.22 at 2kHz, 0.18 at 4 kHz, and 0.1 at 8kHz. These values agreed well with the results from (Goode et al. 1994), which were measured at 114 dB SPL, and also lay near the lower boundary of the experimental results from (O’Connell-Rodwell et al. 2024), measured at 110 dB SPL. These values are lower than the one from the anatomical lever ratio, suggesting some degree of transmission loss possibly from slippage across the IMJ and ISJ, as well as potential rotation and bending of the ossicles, particularly at higher frequencies where three-dimensional motions of the ossicles may become more dominant and these ratios decreased more significantly. As the stimulus level goes up to 170 and 180 dB SPL, our model showed decreased overall SFP-to-umbo transfer ratios to about 0.46 and 0.39 below 1kHz, suggesting compression of transmission at higher levels.

Figures 9b and 9c show incus-to-umbo and SFP-to-incus ratios, which were approximately 0.64 and 0.78, respectively, up to 1 kHz at the stimulus level up to 160 dB SPL. Above 160 dB SPL, these ratios decreased below 1 kHz, consistent with compression of transmission. However, at frequencies above 1 kHz, these ratios did not follow consistent trends. As shown in Fig. 9b, the incus-to-umbo ratio (IMJ slippage) showed a peak at 2 kHz, with values of approximately 0.8 for most SPLs and close to 1.0 at 160 dB SPL. Afterward, this transfer ratio decreased significantly at 4 kHz and remained almost constant up to 8 kHz. In contrast, the SFP-to-incus ratio (ISJ slippage) decreased significantly at 2 kHz, reaching values as low as 0.3 for most levels and 0.07 at 160 dB SPL, as shown in Fig. 9c. At higher frequencies, this ratio showed a local peak at 4 kHz (0.49–0.64, depending on the SPL) and decreased again at 8 kHz. Despite these different trends, the overall SFP-to-umbo transfer ratio (Fig. 9a) demonstrated a much smoother decrease with increasing frequency and SPL.

### 3.7. Spectral Distortions

Although the above-mentioned results present the steady-state response of the model obtained from the most periodic portion of the corresponding signals (i.e., the second half of the time-domain responses), they only represent the fundamental amplitudes. However, spectral distortions, including harmonics and subharmonics, were observed, particularly at high frequencies and high levels. Figure 10a shows the TSD percentages for the ear canal pressure measured near the umbo. For the ear canal pressure, the TSD was small in most cases, with values below 1% up to 0.5 kHz and below 10% up to 2 kHz. Distortions became more noticeable only at 4 and 8 kHz, where the TSD ranged from 16% to 32% at 4 kHz and from 12% to 39% at 8 kHz, depending on the pressure level.

**Fig. 10.**
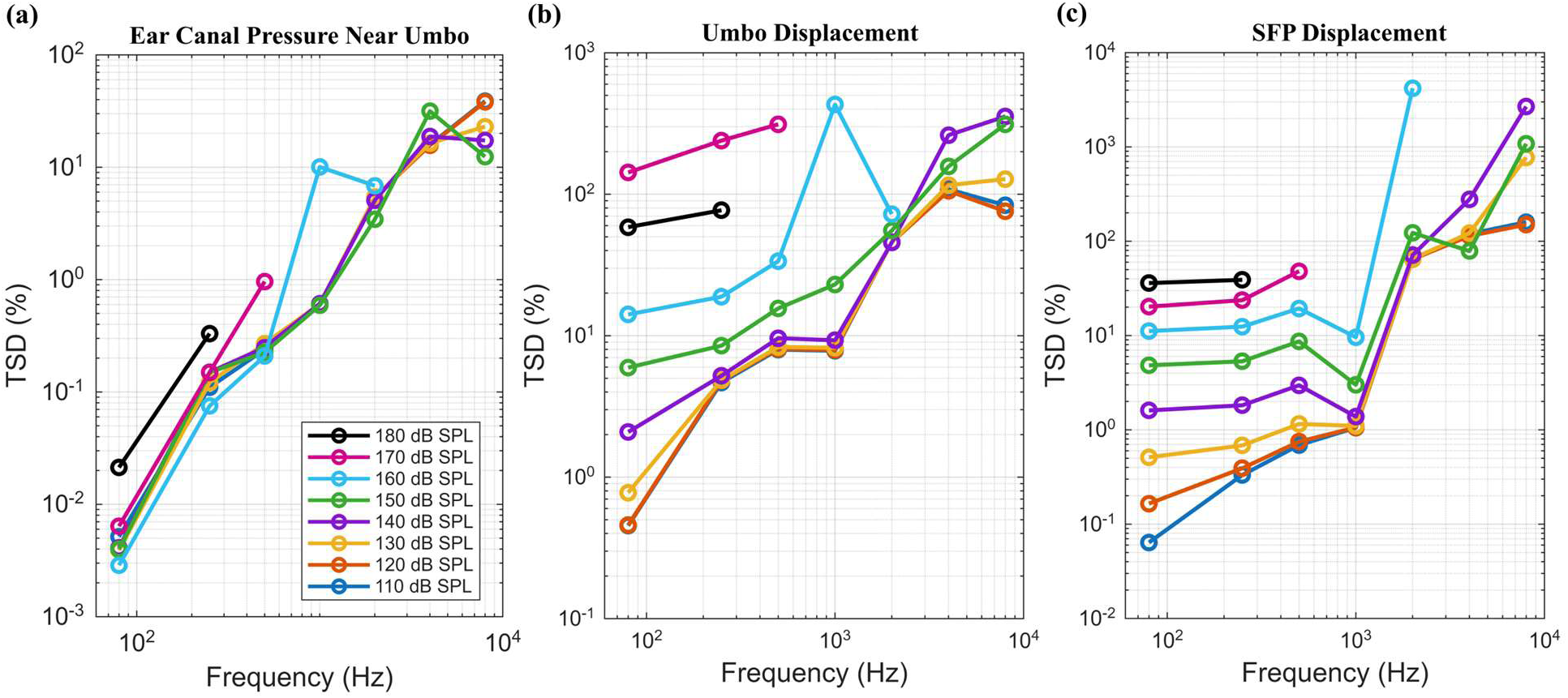
Total spectral distortion (TSD) for the ear canal pressure (a), umbo displacement (b), and SFP displacement (c), computed from the FFT of the second half of the time-domain responses for all investigated frequencies and SPLs. The TSD of the ear canal pressure is negligible in most cases, with only minor distortions observed at the highest frequencies. For the umbo and SFP displacements, the distortions remain negligible or minor for frequencies up to 1 kHz and levels up to approximately 150 dB SPL. At higher frequencies (≥2 kHz) or higher levels (≥160 dB SPL), the distortions contribute substantially to the model responses

Figures 10b and 10c show the TSD values for the umbo (middle) and SFP (right) displacement magnitudes. For the umbo, the TSD was negligible for frequencies up to 1 kHz and levels up to 140 dB SPL, remaining below or around 10%. However, at higher levels (≥150 dB SPL) or higher frequencies (≥2 kHz), the distortion increased substantially. The most extreme cases occurred near the resonance frequency at high levels (i.e., 430% at 1 kHz and 160 dB SPL) and at high-frequency, high-level conditions (354% and 312% at 8 kHz for 140 and 150 dB SPL, respectively). For the SFP, the TSD was small for frequencies up to 1 kHz and levels up to 150 dB SPL, remaining below or around 10%. At higher levels within the 1-kHz condition, the TSD remained below 50%. However, at higher frequencies (≥2 kHz), the TSD increased substantially.

### 3.8. Incremental Nonlinearity of the Umbo and the Stapes Footplate

Figure 11 shows the incremental nonlinearity index in decibel scale for both the umbo (a) and SFP (b), obtained from displacement magnitudes. Blue and red colors indicate compressive (*γ*_dB_ < −0.5 dB) and expansive (*γ*_dB_ > 0.5 dB) nonlinearities, respectively. For umbo at low frequencies (80 to 500 Hz), displacement increased almost linearly (i.e., |*γ*_dB_| ≤ 0.5 dB) with pressure up to 160 dB SPL. Beyond this level, the response exhibited compressive nonlinearity, with indices ranging from −2.9 to −1.9 dB at 170 dB SPL and from −4.9 to −4.7 dB at 180 dB SPL. At higher frequencies, the nonlinearity was initiated at lower stimulus levels. At 1 kHz, the umbo response became nonlinear at 160 dB SPL with an index of −1.8 dB. At 2 kHz, the nonlinearity began at 150 dB SPL with a value of −0.7 dB and increased to −3.2 dB at 170 dB SPL. At 4 and 8 kHz, the umbo displacement exhibited a more complex pattern involving both compressive and expansive nonlinearities. The incremental nonlinearity index ranged from −0.8 to 1.9 dB at 4 kHz and from −3.9 to 0.6 dB at 8 kHz, depending on the sound pressure level.

**Fig. 11.**
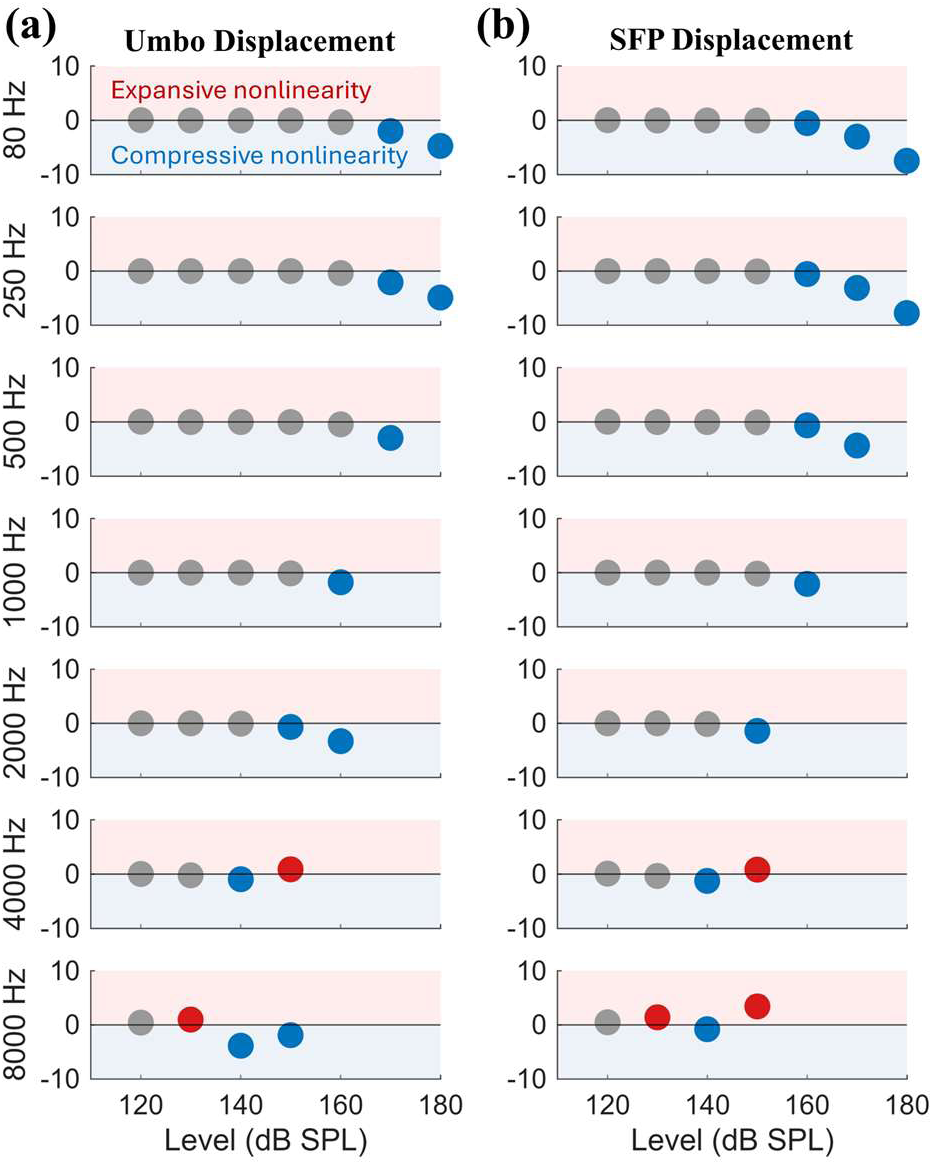
Incremental nonlinearity index in decibels for the umbo (a) and SFP (b) at frequencies ranging from 80 Hz to 8 kHz and excitation levels ranging from 120 to 180 dB SPL. Compressive nonlinearities (γdB < −0.5 dB) and expansive nonlinearities (γdB > 0.5 dB) are shown in blue and red, respectively, while gray points indicate linear or near-linear displacement variation with respect to ear canal pressure. Overall, predominantly compressive nonlinearities were observed for both the umbo and SFP responses at levels around or above 150 dB SPL within the frequency range up to 2 kHz, whereas a combination of compressive and expansive nonlinearities was observed under the 4- and 8-kHz conditions

For the SFP displacement, nonlinear patterns similar to those observed for the umbo displacement were identified. At low frequencies (80–500 Hz), compressive nonlinearity began to appear at 160 dB SPL with values around −0.6 dB and increased to values between −4.4 and −3.0 dB at 170 dB SPL, and between −7.8 and −7.5 dB at 180 dB SPL. At 1 kHz, the nonlinearity began at 160 dB SPL with a value of −2.1 dB, while at 2 kHz it began at 150 dB SPL with a value of −1.4 dB. At 4 kHz, the nonlinearity index ranged from −1.1 to 1.1 dB, whereas at 8 kHz it ranged from −0.9 to 3.5 dB, depending on the sound pressure level. The strongest expansive nonlinearity was observed at 150 dB SPL for both 8 and 4 kHz.

## 4. Discussion

In this study, we developed a nonlinear acoustic–structural FE model of the human middle ear and ear canal for the time-domain analysis. We then analyzed the model’s response to pure-tone sound pressure signals spanning a wide frequency range, from 80 Hz to 8 kHz, under varying sound pressure levels ranging from 110 to 180 dB SPL.

### 4.1. Validation and Comparison with Experimental Data

A comparable range of normalized displacements was observed between the present FE simulations and the experimental measurements reported in the literature (Gan et al. 2004b). However, some discrepancies remained, particularly at low to mid frequencies for the umbo displacement, where the model predicted smaller normalized values than those reported experimentally. It should be noted that the experimental measurements in (Gan et al. 2004b) primarily represent motion in the LM direction and were quantified using peak-to-peak displacements in the time domain. In contrast, the present study evaluates the three-dimensional displacement magnitude based on the fundamental component in the frequency domain. These differences in measurement definitions may contribute to the observed discrepancies. In addition, the sound pressure levels applied in the experimental study differed from those considered in the present simulations, which may have further influenced the comparison.

The FE results also showed good agreement with qualitative mode shapes of the TM and ossicular chain, as well as the three-dimensional full-field displacement magnitudes obtained from synchrotron-based X-ray phase-contrast microtomography at low frequencies (Rodgers et al. 2026). For example, at 120 dB SPL and 512 Hz, the experimentally measured TM displacement magnitudes (peak-to-peak values divided by two) ranged from nearly zero in regions close to the annulus to approximately 3 μm in the PS quadrant located midway between the umbo and annulus, with a displacement magnitude of approximately 1 μm at the umbo. These values are consistent with our FE results at 120 dB SPL and 500 Hz. Nevertheless, the three-dimensional geometry and the Ogden material parameters used in the present work were derived from different experimental datasets, introducing potential inter-subject variability in comparison with the experimental measurements.

### 4.2. Ear Canal Acoustic Response and Middle-Ear-Induced Distortions

At low to mid frequencies (80 Hz to 4 kHz), the acoustic pressure near the umbo was nearly equal to that prescribed at the input plane of the ear canal, indicating a spatially uniform acoustic field due to the relatively long wavelength of sound at these frequencies compared to the canal length. At higher frequencies (e.g., 8 kHz), the pressure near the umbo increased compared to that at the entrance. As the wavelength (approximately 4 cm at 8 kHz) becomes comparable to the dimensions of the modeled ear-canal segment, spatial pressure variations develop along the canal, resulting in increased pressure near the umbo (Ravicz et al. 2007, 2014, 2019; Motallebzadeh et al. 2017).

The pressure signals across all levels and frequencies exhibited a nearly pure-tone behavior. However, at higher frequencies, the responses became slightly distorted compared to a perfect sinusoidal waveform, with some non-harmonic spectral components appearing in the frequency-domain representations. Although the acoustic domain is governed by linear wave propagation, these nonlinear features arise from coupling with the TM and middle-ear structures, which exhibit level-dependent nonlinear responses. Consequently, nonlinear distortion motions within the structural domain of the TM generate pressures that propagate in the reverse direction back into the ear canal as reflections. This effect became more pronounced at higher frequencies, where structural dynamics and coupling effects played increasingly significant roles.

### 4.3. Modal Transition and Nonlinear Dynamic Response of the TM and Ossicular Chain

The spatial vibration patterns of the TM and ossicular chain exhibited a clear frequency-dependent transition from simple to complex mode shapes (Fig. 2). At the representative low frequency (0.5 kHz), the TM response was dominated by a low-order mode characterized by a relatively smooth and continuous displacement distribution, indicating near-uniform TM motion. The ossicular chain at this frequency also demonstrated coherent motion dominated by a primary mode. However, due to the presence of joints, ligaments, and tendon attachments, regions near structural constraints exhibited reduced displacement (not shown), resulting in a constrained yet coordinated motion pattern. Other than a slight increase in the anterior portion of the TM relative to the posterior portion, the responses generally appear to scale with level.

At mid frequency (4 kHz), the TM response became more spatially structured, with multiple localized regions of high displacement emerging across the TM surface, indicating the activation of higher-order vibration modes. Correspondingly, the ossicular chain exhibited increased spatial variation in displacement, reflecting the combined effects of joint flexibility, ligament constraints, and inertial contributions. The motion was no longer confined to a single rotational axis and included multi-axis (rocking) components. For additional details, see the animations of TM and ossicular-chain motion and the three-dimensional trajectories of umbo and SFP motion at representative frequencies and sound pressure levels provided in the SM.

At high frequency (8 kHz), the TM displayed even more non-uniform and spatially distributed vibration patterns, characterized by multiple localized displacement peaks consistent with a multi-modal response. Similarly, the ossicular chain exhibited complex motion with substantial spatial variability, indicating the contribution of multiple interacting vibration modes that arose naturally from the full-field FE solution rather than from *a priori* selection of specific modes. The displacement trajectories of both the umbo and SFP further reflected this transition, evolving from simple, nearly one-dimensional piston-like motion at low frequency to two-dimensional planar and fully three-dimensional multi-directional trajectories at higher frequencies (Fig. 2c and d).

At 8 kHz, the temporal evolution of the Poincaré trajectories also differed between excitation levels. At 120 dB SPL, the trajectory expanded over time, indicating an increase in the effective displacement at the measurement location. In contrast, at higher input levels, later samples exhibited reduced amplitudes compared to earlier samples. This behavior is attributed to the complex spatial vibration patterns of the TM and other middle-ear structures at high frequencies, where multiple local displacement maxima with different phase relationships coexist. At lower excitation levels, the modal contributions remained relatively phase-aligned, leading to constructive superposition and amplitude growth. This is consistent with the summation of different TM modes hypothesized by (Fay et al. 2006). However, at higher levels, nonlinear effects altered the modal phase relationships and promoted inter-modal coupling, resulting in partial destructive interference and a reduction in the measured displacement amplitude over time.

### 4.4. DC Shift and Asymmetric Response under Large Deformation

We observed a DC shift in the umbo displacement (and more generally across the entire TM surface) as well as in the SFP displacement under high-frequency and high-level excitation conditions. This characteristic may be associated with geometric nonlinearity under large deformation. For example, a gradual outward (lateral) shift of the TM was observed at 8 kHz and 150 dB SPL. Because the TM possesses an initially curved conical geometry, inward (medial) deformation (due to a positive ear canal pressure) toward the middle-ear cavity produces greater stretching and stress stiffening on the TM than outward lateral deformation toward the ear canal due to a negative ear canal pressure. Consequently, the instantaneous restoring stiffness during inward motion becomes larger than that during outward motion, resulting in an asymmetric oscillatory response and a net lateral shift in the mean position (Radenković et al. 2015; Cattiaux et al. 2020; Mohammadi et al. 2024c). The DC shift observed at the SFP may similarly result from transmission of the asymmetric TM/umbo response through the ossicular chain, with additional modulation by nonlinear deformation of the ossicular joints and SAL. This effect is further amplified at high frequencies due to nonlinear modal interactions and multi-axis ossicular motion under strongly nonlinear vibration conditions.

### 4.5. Nonlinear Distortions at High Frequencies and High Sound Levels

The TSD results revealed substantial distortions at high frequencies (≥2 kHz) and high stimulus levels (≥160 dB SPL). These distortions manifested as amplitude modulation, beating-envelope patterns, and other deviations from purely sinusoidal responses, affecting both the waveform shape and the effective range of motion in the time domain. Consequently, the steady-state responses derived from the frequency-domain analyses under these high-frequency, high-level conditions should be interpreted with caution. In such cases, the amplitudes obtained from the fundamental component represent prescribed displacement magnitudes rather than the true peak displacements observed in the time-domain signals.

### 4.6. Transient Response and Steady-State Convergence

The normalized Euclidean difference between adjacent temporal segments generally increased with frequency; however, a pronounced peak was observed around 2 kHz. This behavior corresponds to the transition region just above the middle-ear resonance, where the system exhibits strong phase sensitivity and interference effects. In this regime, small phase variations between successive cycles can lead to disproportionately large differences in the time-domain signals, resulting in elevated Euclidean difference values. This observation is consistent with previous stochastic analyses reporting maximum uncertainty amplification and phase variability in the 2–3 kHz range (Ebrahimian et al. 2023b). For frequencies up to 1 kHz, the middle-ear responses stabilized rapidly, typically within two or three cycles, indicating a short transient stage. This finding suggests that fewer cycles are sufficient for low-frequency analyses in future simulations, reducing computational costs.

### 4.7. Transfer Ratios Across the Ossicular Chain

The SFP-to-umbo transfer ratio at frequencies up to 1 kHz and levels up to 160 dB SPL was around 0.5, which lies within the range reported in experimental measurements. For instance, an approximate velocity ratio of 0.5–0.7 for frequencies up to 1 kHz was reported by (O’Connell-Rodwell et al. 2024). Their measurements also showed that the SFP-to-umbo ratio decreased at higher frequencies to values as low as approximately 0.1, which is similar to the computational results obtained in the present study. Furthermore, the low-frequency incus-to-umbo transfer ratio of approximately 0.64 is reasonably consistent with the classical malleus-incus lever ratio of approximately 1.25–1.6 reported in the literature (Péus et al. 2020; Schmeltz et al. 2024), which would correspond to an idealized incus-to-umbo displacement ratio of approximately 0.63–0.8 under rigid-body lever motion.

An interesting observation was the different frequency-dependent dynamic behaviors observed in different parts of the ossicular chain. At low frequencies, the malleus, incus, and stapes moved almost as a single unit, resulting in nearly constant transfer ratios. As the sound level increased, the contribution of the nonlinear Ogden material model led to lower transfer ratios, particularly at the highest investigated levels (160–180 dB SPL). At higher frequencies, however, opposite trends were observed in the incus-to-umbo and SFP-to-incus transfer ratios. At 2 kHz, the incus-to-umbo ratio exhibited a local peak while the SFP-to-incus ratio showed a local trough, indicating relatively efficient transmission from the umbo to the incus but reduced transmission from the incus to the stapes. At 4 kHz, the opposite behavior was observed. These frequency-dependent variations likely reflect changes in ossicular dynamics, including bending and relative motion at the ossicular joints, which become increasingly important at higher frequencies as the motion transitions away from rigid-body behavior. Because the two transfer ratios exhibited complementary trends, their product, i.e., the overall SFP-to-umbo transfer ratio, showed a much smoother decrease with increasing frequency and SPL despite the substantial variations observed in the individual ratios.

### 4.8. Compressive and Expansive Nonlinearity in Middle-Ear Response

The proposed nonlinear middle-ear model, employing a first-order Ogden hyperelastic formulation, predominantly exhibited compressive nonlinearity in the umbo and SFP displacements with increasing canal pressure. Compressive nonlinearity was more prominent in SFP than in the umbo, which is aligned with the experimental observation (Cheng et al. 2021). Such compressive nonlinearity at high sound levels (typically above 130 dB SPL) has been reported in the literature (Guinan and Peake 1967; Cheng et al. 2021) and is often associated with protective mechanisms that limit the transmission of excessive sound energy to the inner ear (Price and Kalb 1991). However, expansive nonlinearity has also been reported in some specimens and frequency ranges (Cheng et al. 2021; Rodgers et al. 2026). In the present model, the onset of nonlinear behavior at low frequencies generally occurred at higher stimulus levels (above 150 dB SPL) compared to experimental reports where nonlinear responses were observed at approximately 110-150 dB SPL, depending on frequency and measurement location (Cheng et al. 2021). The discrepancy may partly reflect uncertainties in the constitutive properties used for the soft tissues. The Ogden parameters were obtained from separate experimental studies performed on different specimens and, in some cases, under loading and boundary conditions that differed substantially from the intact middle-ear configuration. Consequently, the effective *in situ* mechanical behavior of these tissues within the complete middle-ear system may differ from the material properties used in the model.

The observed predominance of compressive nonlinearity in the model can be partly attributed to the Ogden material parameters used, particularly the relatively large values of the exponent α (Table 1). Larger α values promote stronger strain-stiffening responses, leading to an increase in effective stiffness with deformation amplitude, which in turn contributes to sublinear (compressive) growth of displacement with respect to input loading. In contrast, values of α closer to 2 result in a more nearly linear response, while smaller values may lead to a more compliant response and potentially expansive nonlinearity under certain conditions (Lohr et al. 2022). In addition to material nonlinearity, geometric nonlinearity also plays an important role. Large-deformation effects, including the use of the Green–Lagrange strain measure and deformation-dependent structural stiffness, introduce amplitude-dependent behavior (Li and Zhan 2000). Furthermore, the complex geometry and coupled dynamics of the middle ear contribute to nonlinear dynamics through modal interactions and multi-directional motion (Karamooz Mahdiabadi et al. 2021).

### 4.9. Limitations and Future Directions

One limitation of the proposed model is that we only investigated a first-order Ogden hyperelastic formulation for the soft tissues of the middle ear, while the potential capabilities of other hyperelastic models in middle-ear simulations remain largely unknown, mainly due to the lack of experimentally identified material parameters for middle-ear structures in the literature. Also, the Ogden constants reported in the literature come from different types of experimental measurements (usually pure tension or pure shear studies), use different constitutive formulations than the one implemented in COMSOL, and are identified under different loading conditions than those applied in our model which includes a combination of bending, tension, compression, pressure, and multiaxial stretching. In addition, the Ogden parameters used in the current study were derived from a limited number of *ex vivo* experimental measurements on a small set of specimens and therefore may not fully represent the inter-individual variability in humans. Accordingly, the influence of variations in higher-order Ogden nonlinearities on middle-ear responses remains to be investigated.

Furthermore, the current model did not incorporate the anisotropic properties of the TM (Fay et al. 2005). A potential improvement for future studies would be the implementation of the Holzapfel–Gasser–Ogden (HGO) model (Holzapfel et al. 2004; Nolan et al. 2014), which incorporates fiber orientation and could better capture the anisotropic behavior of the TM arising from its radial and circumferential collagen fibers. In addition, several studies have suggested the presence of prestrain and prestress in the TM (Muyshondt and Dirckx 2022; Livens and Dirckx 2023), which were neglected in the present study.

The presented nonlinear framework provides a computational tool for investigating middle-ear mechanics under high-intensity acoustic excitation. In particular, the model can support future studies on nonlinear sound transmission through the middle ear and its potential role in acoustic trauma and hearing-loss mechanisms under extreme sound pressure levels. The ability of the model to capture time-domain nonlinear responses over a wide range of frequencies and sound pressure levels also contributes to the evaluation of middle-ear protective mechanisms and the development of improved hearing-protection strategies for high-noise environments, such as in industrial factories.

Moreover, the framework may serve as a basis for future coupled middle-ear/cochlear models aimed at investigating how nonlinear middle-ear behavior influences intracochlear pressure transmission under intense acoustic loading. Furthermore, improving the understanding of the nonlinear mechanical responses of middle-ear soft tissues can support future tissue-engineering and regenerative-medicine approaches, including the design of biomimetic scaffolds and engineered graft materials for repair and reconstruction of the TM and other middle-ear tissues (Trakimas et al. 2018; Kozin et al. 2019; Ghanad et al. 2021).

## 5. Conclusions

A nonlinear time-domain finite-element model of the human middle ear with the Ogden hyperelastic formulation was developed to study the nonlinear behaviors of the middle ear under high-intensity acoustic excitation. The three-dimensional displacement patterns of the TM and ossicular chain at low frequencies, for which experimental full-field measurements are available, were consistent with previously reported observations. In addition, the normalized displacement magnitudes of the umbo and SFP showed good agreement with experimental measurements reported in the literature. The proposed FE model enabled the analysis of transient and steady-state full-field three-dimensional middle-ear displacements over a wide range of frequencies and sound pressure levels, including very high levels up to 180 dB SPL. The displacement trajectories and full-field vibration patterns revealed a frequency-dependent transition from predominantly piston-like motion at frequencies up to 1 kHz (below the resonance frequency) to more complex multi-axis and rocking motion at frequencies ≥2 kHz (above the resonance frequency). Nonlinear behavior appeared in several forms, including harmonic generation at high stimulus levels (>150 dB SPL), subharmonics and modulated spectral distortions at high frequencies (≥2 kHz), and compressive (maximum nonlinearity index of −7.8 dB at 180 dB SPL for SFP) and expansive (maximum index of +3.4 dB at 8 kHz for SFP) displacement responses with increasing sound pressure level. The observed nonlinear responses arise from the combined effects of material nonlinearity, geometric nonlinearity associated with large deformation, and dynamic structural interactions, including modal coupling and multi-axis motion of the ossicular chain. Time-domain analyses further showed that the steady-state response was generally established within the first three cycles for frequencies up to 1 kHz, whereas longer transient times were observed at higher frequencies.

## Acknowledgments

This work was supported by the National Institutes of Health (NIH), National Institute on Deafness and Other Communication Disorders (NIDCD), grant number R01 DC016079 to JTC. We gratefully acknowledge Daniel Cardosi for valuable discussions on nonlinear time-series analysis.

## Author contributions

H.M.: Writing—original draft, Conceptualization, Methodology, Investigation, Software, Formal analysis, Data curation, Visualization. A.T.: Writing—review and editing, Methodology, Software, Investigation, Visualization. S.P.: Writing—review and editing, Conceptualization, Methodology, Supervision. J.T.C.: Writing—review and editing, Conceptualization, Methodology, Supervision, Funding acquisition.

## Supplementary materials

Supplementary materials related to this article can be found in the online version.

## Data availability

The COMSOL models and data generated and/or analysed during the current study are available via Harvard Dataverse at https://dataverse.harvard.edu/dataverse/Otobiomech.

## ^1^List of abbreviations

AML: anterior malleal ligament
AP: anterior–posterior
FE: finite-element
FFT: fast Fourier transform
HGO: Holzapfel–Gasser–Ogden
IMJ: incudomalleolar joint
ISJ: incudostapedial joint
LM: lateral–medial
LML: lateral malleal ligament
LP: lenticular process
PIL: posterior incudal ligament
PS: posterior–superior
SAL: stapedial annular ligament
SFP: stapes footplate
SI: superior–inferior
SIL: superior incudal ligament
SM: Supplementary Materials
SML: superior malleal Ligament
SPL: sound pressure level
ST: stapedial tendon
TM: tympanic membrane
TSD: total spectral distortion
TTT: tensor tympani tendon

## Footnotes

2 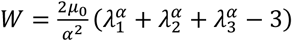 where 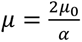

