## Supplementary Materials for "A nonlinear time-domain finite-element model of the human middle ear subjected to intense sound waves"

### Slide 1
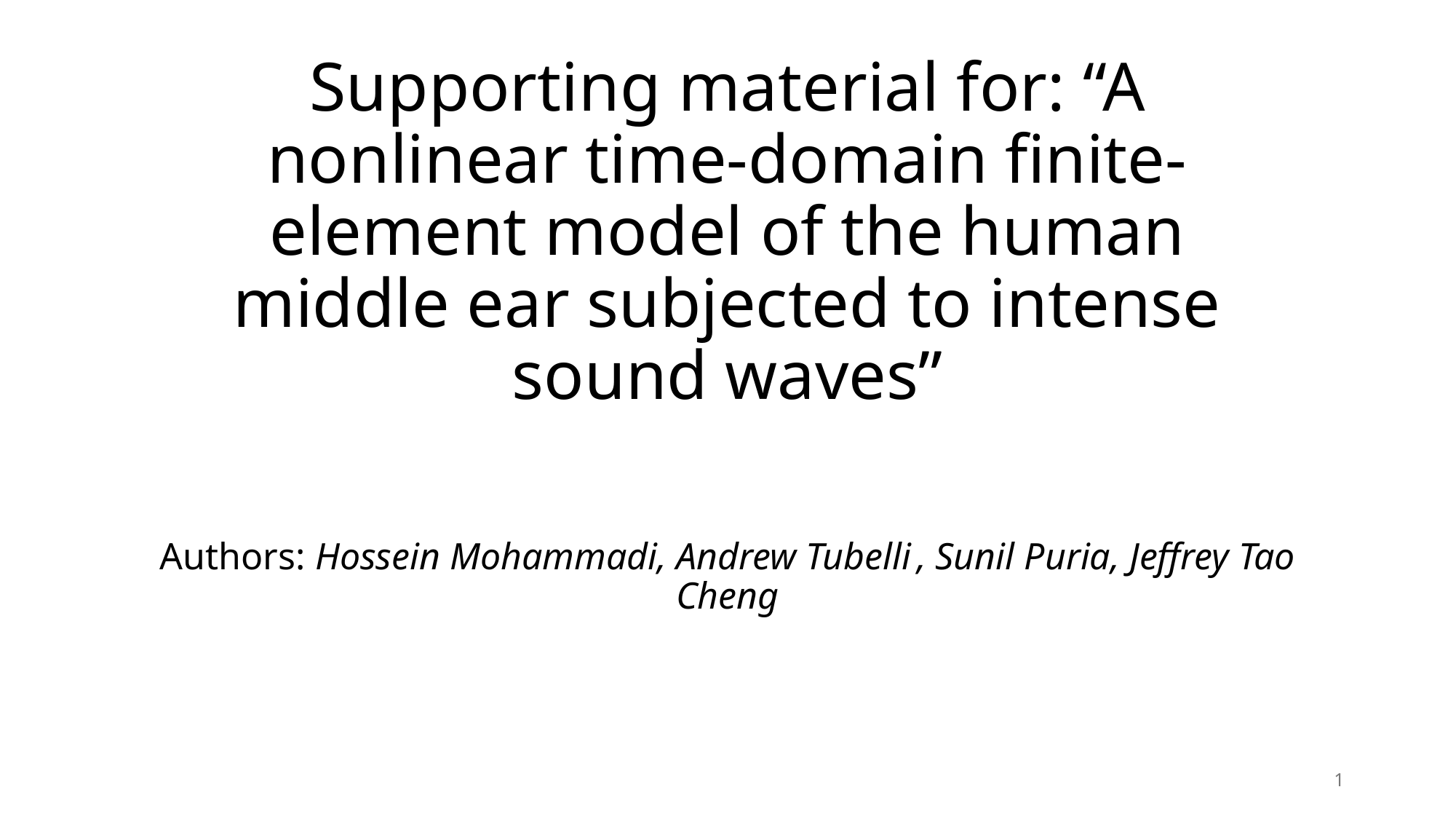

#### Supporting material for: “A nonlinear time-domain finite-element model of the human middle ear subjected to intense sound waves”
Authors: Hossein Mohammadi, Andrew Tubelli , Sunil Puria, Jeffrey Tao Cheng
1

### Slide 2
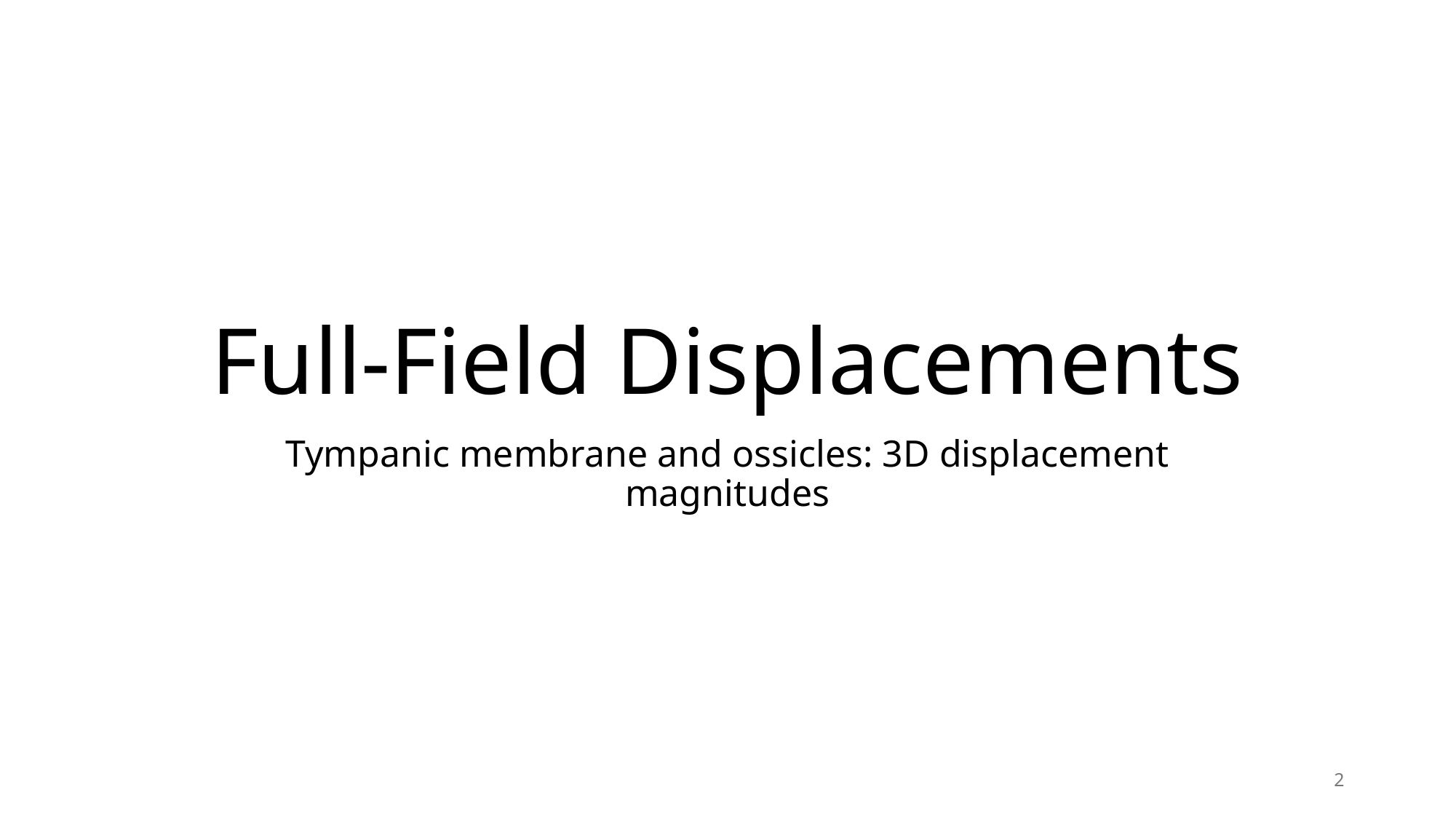

#### Full-Field Displacements
Tympanic membrane and ossicles: 3D displacement magnitudes
2

### Slide 3
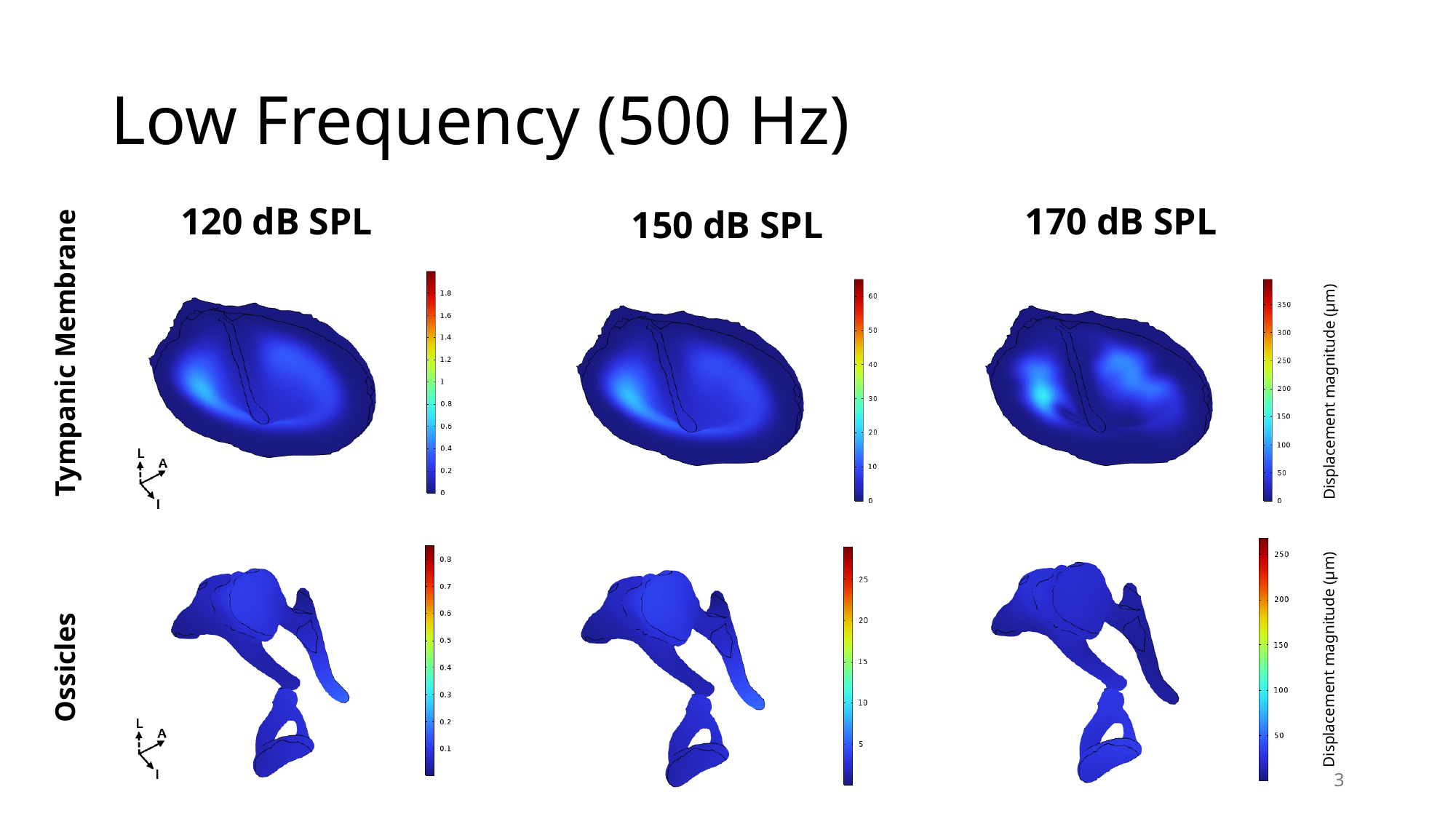

#### Low Frequency (500 Hz)
120 dB SPL
170 dB SPL
150 dB SPL
Tympanic Membrane
Displacement magnitude (µm)
Displacement magnitude (µm)
Ossicles
3

### Slide 4
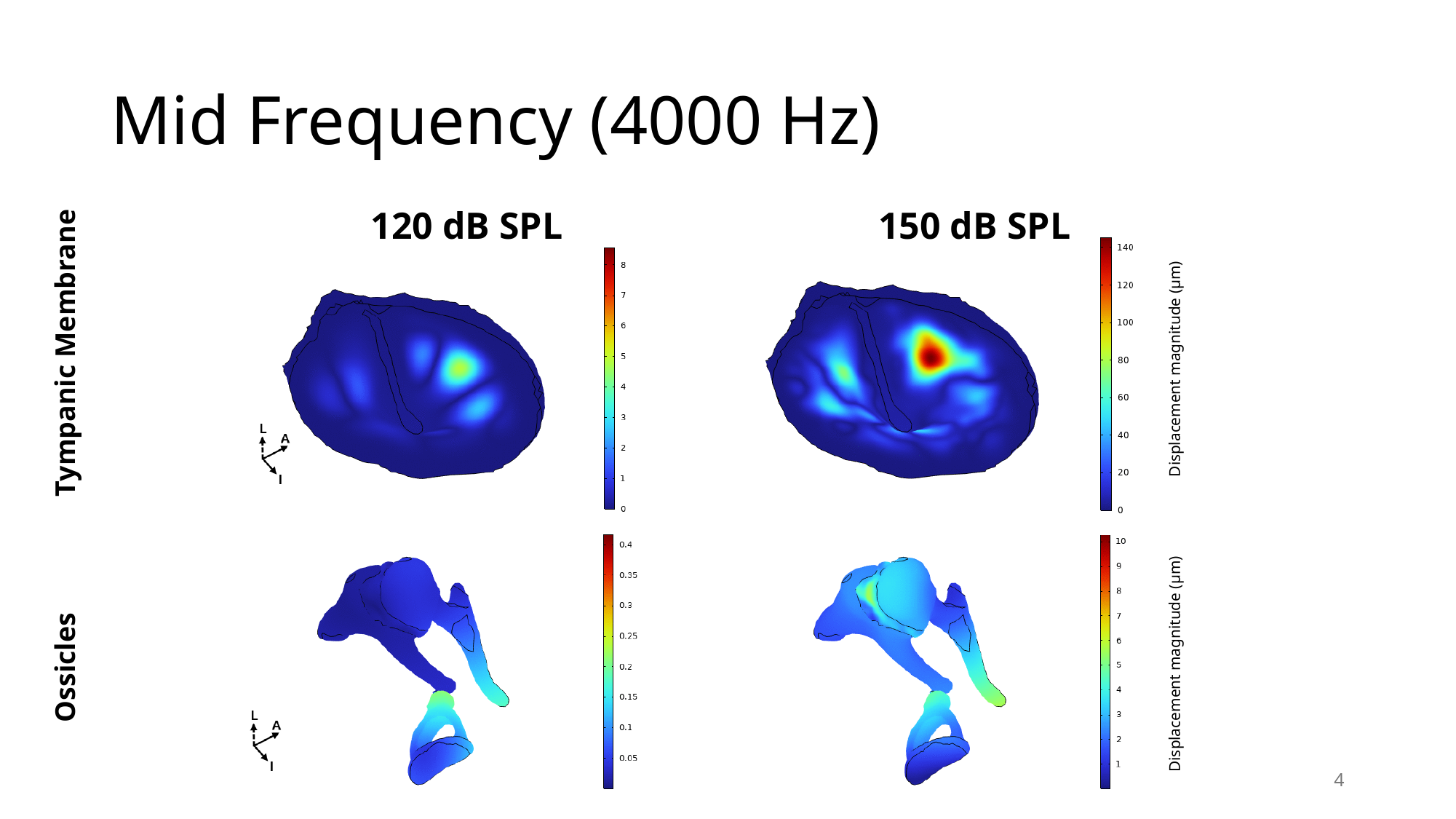

#### Mid Frequency (4000 Hz)
150 dB SPL
120 dB SPL
Tympanic Membrane
Displacement magnitude (µm)
Ossicles
Displacement magnitude (µm)
4

### Slide 5
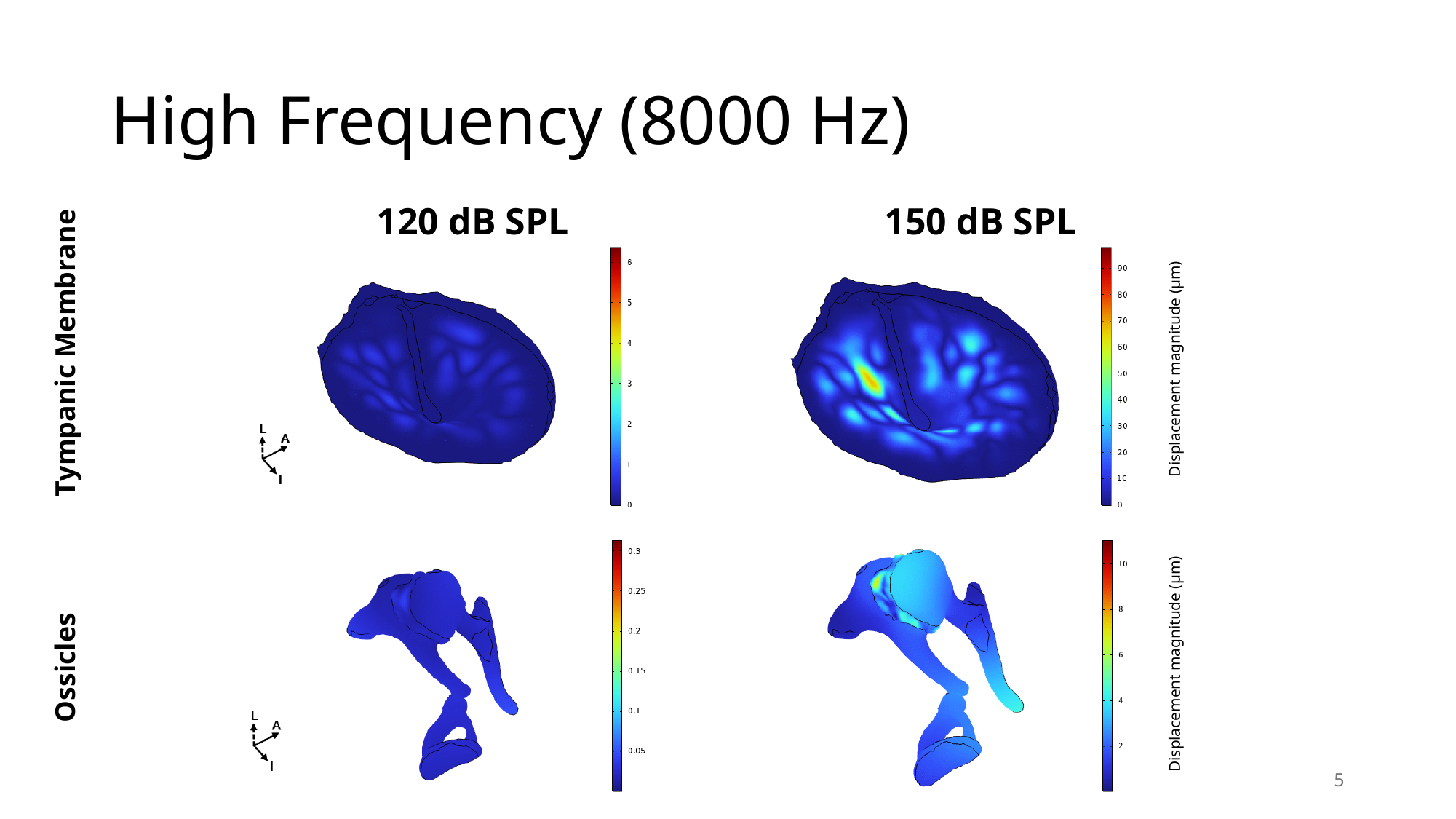

#### High Frequency (8000 Hz)
150 dB SPL
120 dB SPL
Tympanic Membrane
Displacement magnitude (µm)
Ossicles
Displacement magnitude (µm)
5

### Slide 6
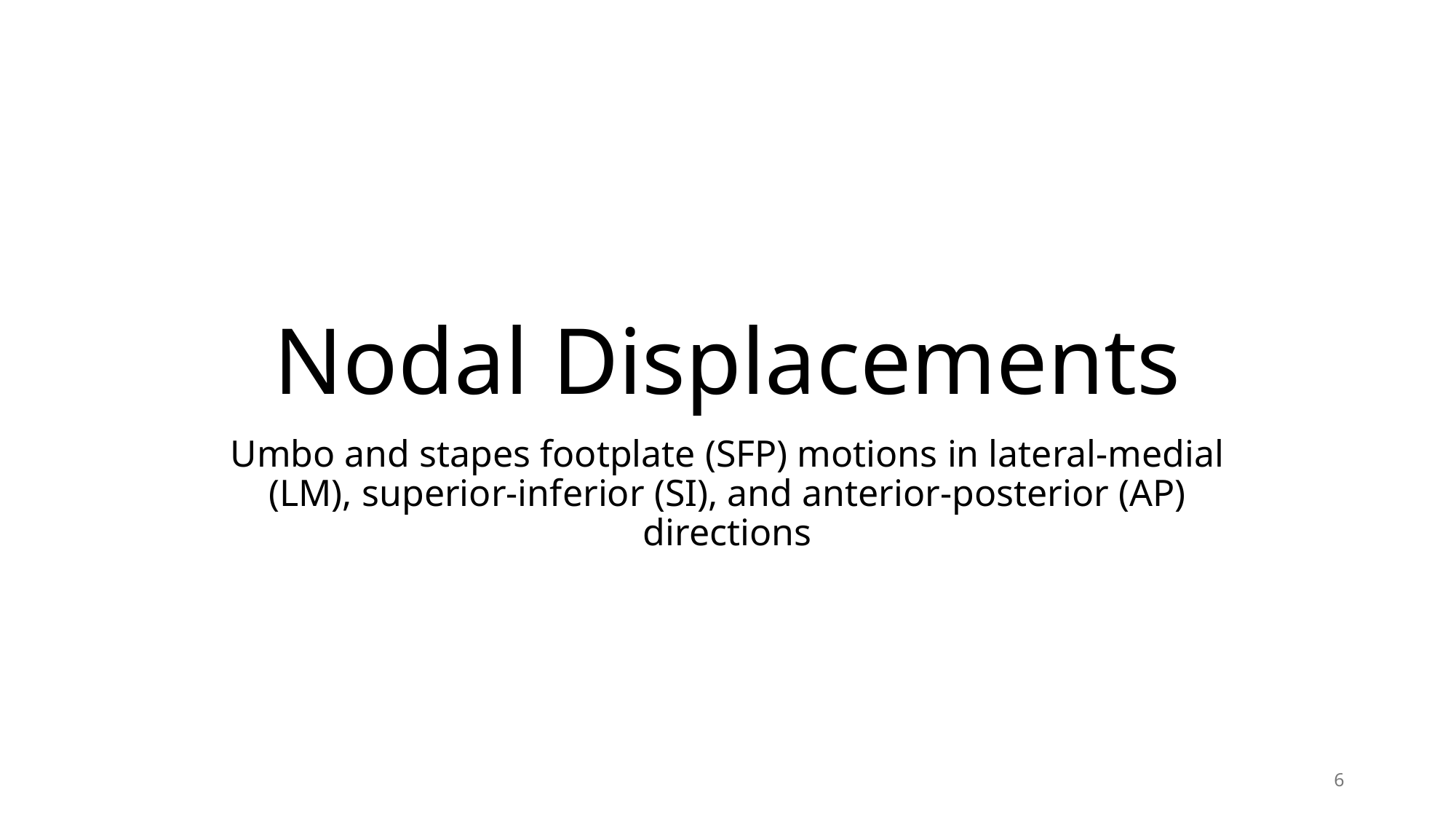

#### Nodal Displacements
Umbo and stapes footplate (SFP) motions in lateral-medial (LM), superior-inferior (SI), and anterior-posterior (AP) directions
6

### Slide 7
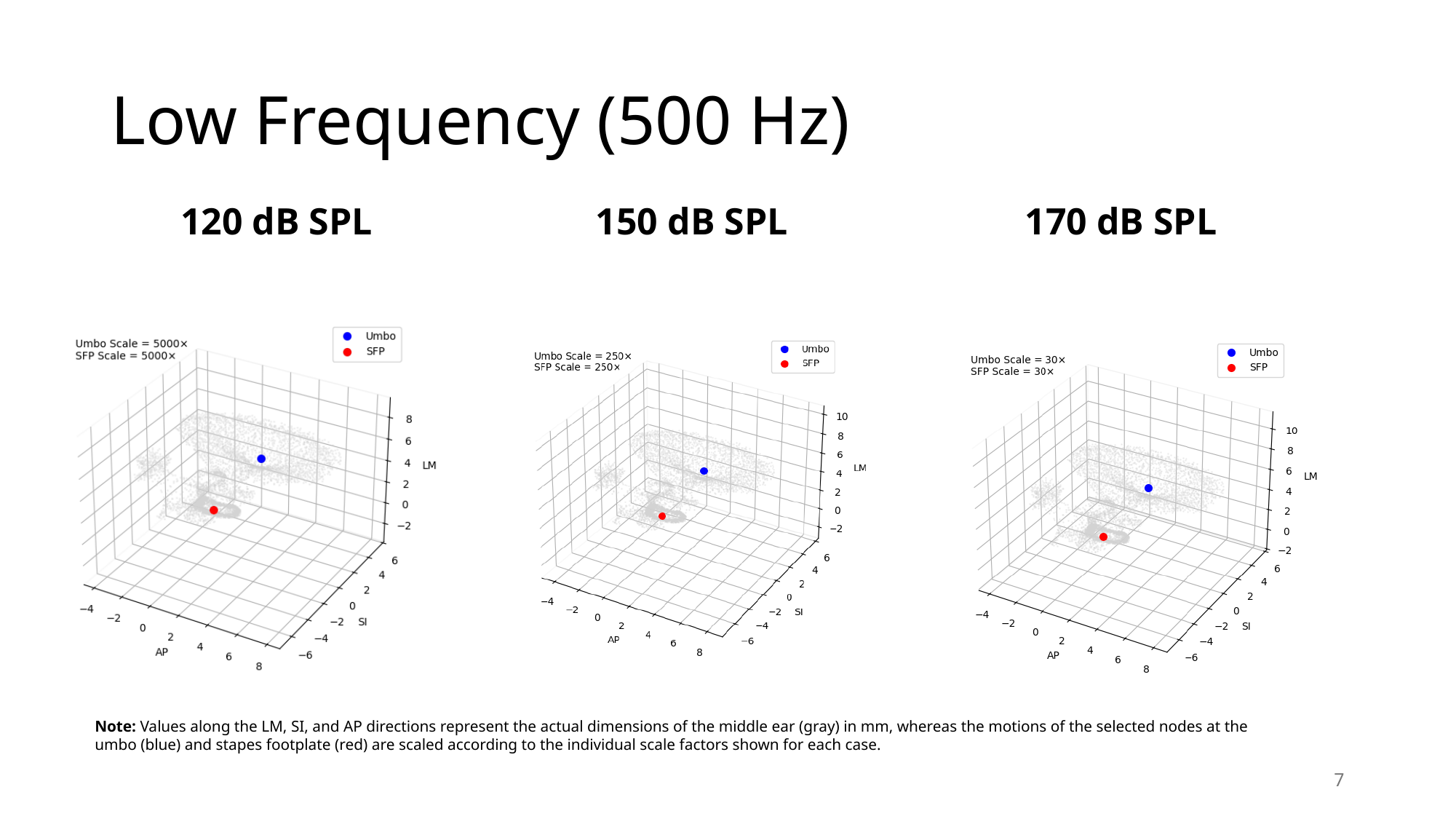

#### Low Frequency (500 Hz)
120 dB SPL
150 dB SPL
170 dB SPL
Note: Values along the LM, SI, and AP directions represent the actual dimensions of the middle ear (gray) in mm, whereas the motions of the selected nodes at the umbo (blue) and stapes footplate (red) are scaled according to the individual scale factors shown for each case.
7

### Slide 8
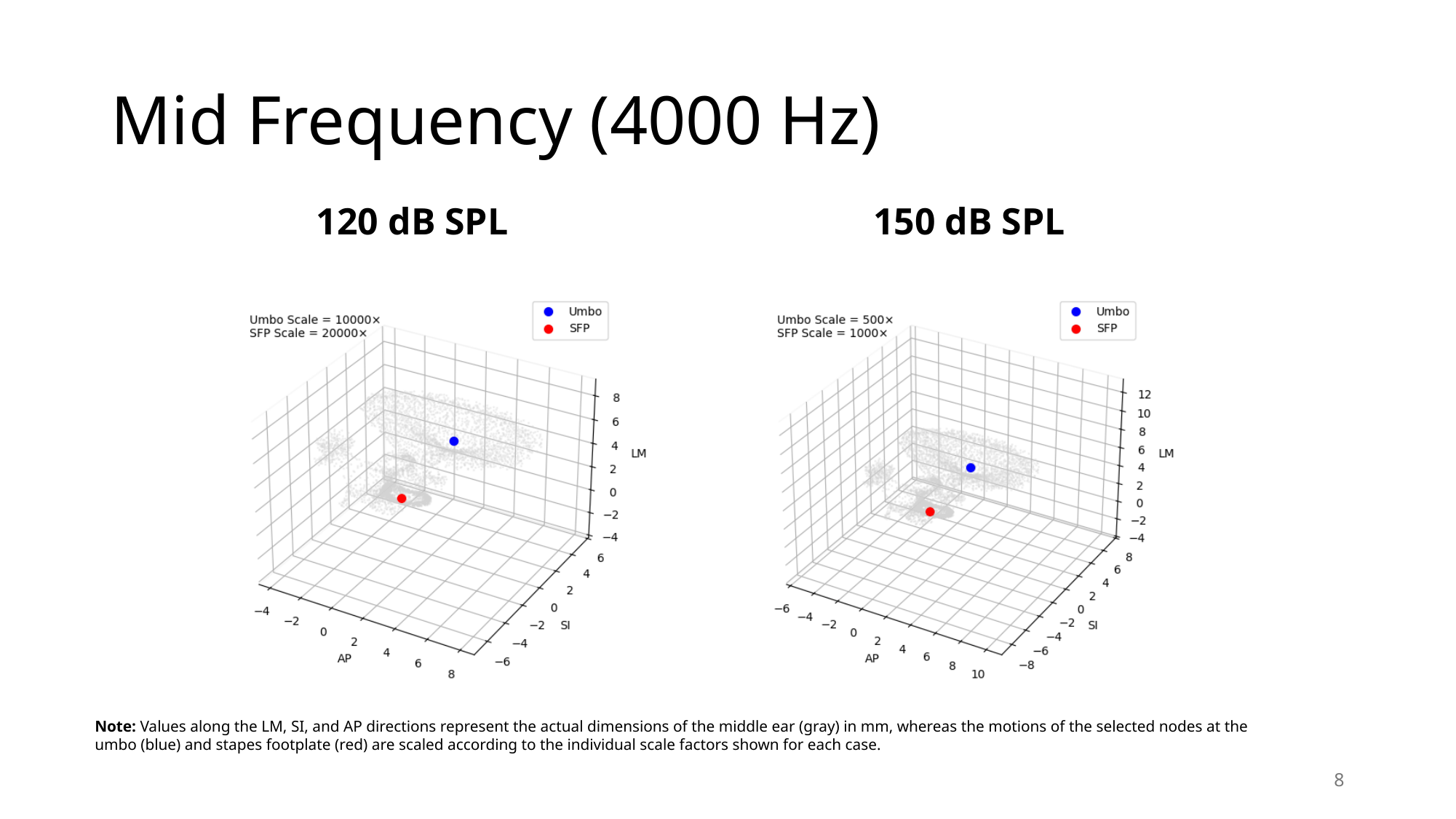

#### Mid Frequency (4000 Hz)
120 dB SPL
150 dB SPL
Note: Values along the LM, SI, and AP directions represent the actual dimensions of the middle ear (gray) in mm, whereas the motions of the selected nodes at the umbo (blue) and stapes footplate (red) are scaled according to the individual scale factors shown for each case.
8

### Slide 9
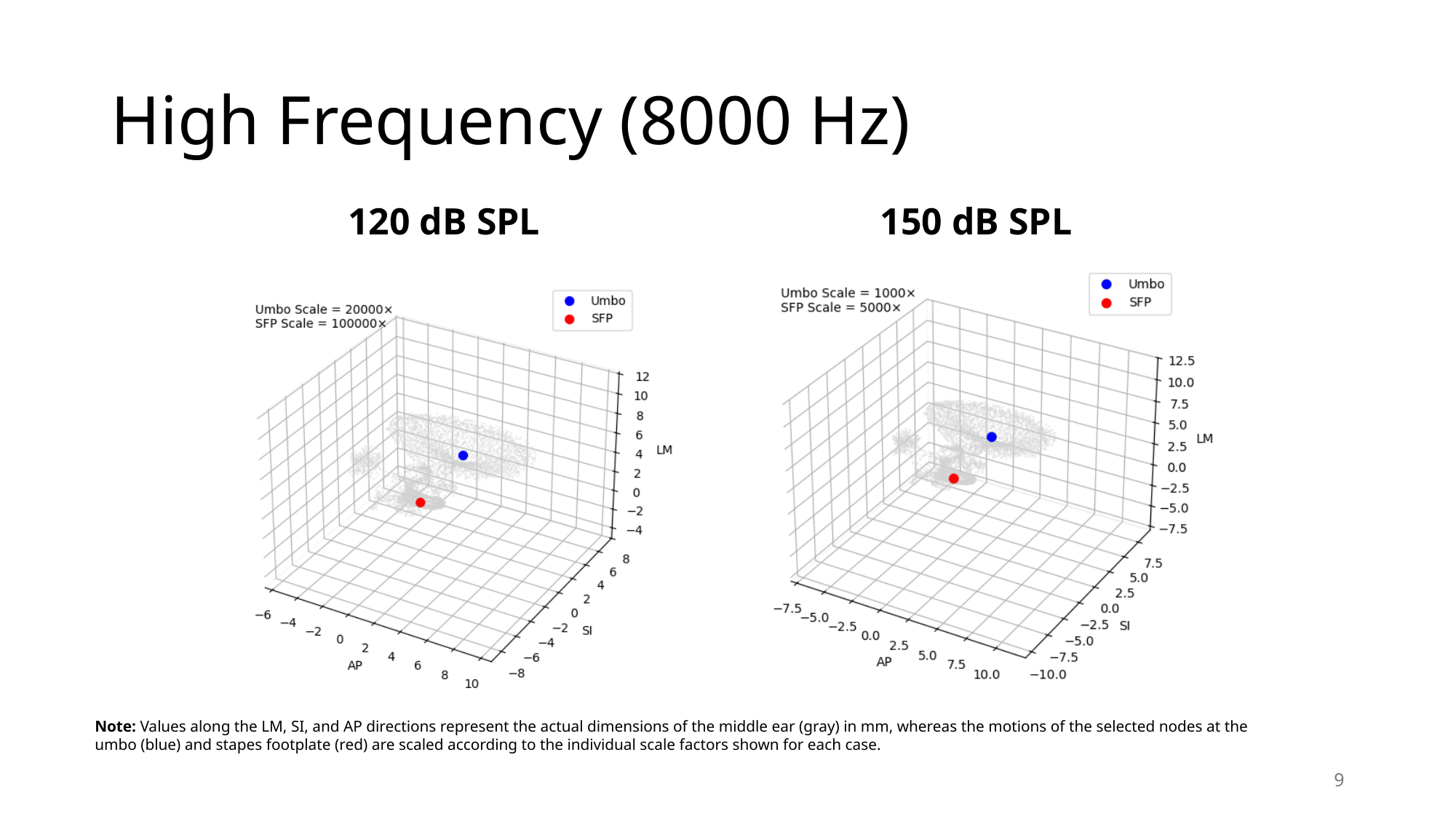

#### High Frequency (8000 Hz)
120 dB SPL
150 dB SPL
Note: Values along the LM, SI, and AP directions represent the actual dimensions of the middle ear (gray) in mm, whereas the motions of the selected nodes at the umbo (blue) and stapes footplate (red) are scaled according to the individual scale factors shown for each case.
9
